# Selection of Synthetic Proteins to Modulate the Human Frataxin Function

**DOI:** 10.1101/2022.02.11.480108

**Authors:** María Florencia Pignataro, María Georgina Herrera, Natalia Fernández, Martín Aran, Fernando Bataglini, Javier Santos

## Abstract

Frataxin is a kinetic activator of the mitochondrial supercomplex for iron–sulfur cluster assembly. Low frataxin expression or a decrease in its functionality results in Friedreich’s Ataxia (FRDA). With the aim of creating new molecular tools to study this metabolic pathway, and ultimately, to explore new therapeutic strategies, we have investigated the possibility of obtaining small proteins exhibiting a high affinity for frataxin. In this study, we applied the ribosome display approach, using human frataxin as the target. We focused on Affi_224, one of the proteins that we were able to select after five selection rounds. We have studied the interaction between both proteins and discussed some applications of this specific molecular tutor, concerning the modulation of supercomplex activity. Affi_224 and frataxin showed a *K*_D_ value in the nanomolar range, as judged by surface plasmon resonance analysis. Most likely, it binds to the frataxin acidic ridge, as suggested by the analysis of chemical shift perturbations (NMR) and computational simulations. Affi_224 was able to increase Cys NFS1 desulfurase activation exerted by the FRDA frataxin variant G130V. Our results suggest quaternary addition may be a new tool to modulate frataxin function *in vivo*. Nevertheless, more functional experiments under physiological conditions should be carried out to evaluate Affi_224 effectiveness in FRDA cell models.

## INTRODUCTION

Human frataxin is a small protein that works as a kinetic activator of the protein supercomplex for iron–sulfur cluster biosynthesis in the mitochondrial matrix, in eukaryotic cells [1–6]. Low frataxin expression or a decrease in its functionality results in Friedreich’s Ataxia (FRDA), a cardio neurodegenerative disease, which is experienced by approximately 1 out of every 50,000 individuals [7, 8].

Frataxin is synthesized as a precursor of 210 residues. It is imported to the mitochondria and processed in two steps [9]. The first step results in an intermediary protein that comprises residues 42 to 210. The second step results in the mature form that comprises residues 81 to 210. The stretch formed by residues 81-92 is disordered, whereas the rest of the protein is structured [10]. Frataxin is a globular protein and its stability is approximately 9 kcal mol^−1^[11, 12]. The structure of the human frataxin has been obtained by crystallography [13] and NMR [14, 15]. The last part of the protein, the C-terminal region (residues 196-210, CTR), is essential for FXN stability [11, 12, 16] and it lacks a periodic structure (**Figures 1A and B**). Remarkably, some frataxin variants, that result in FRDA, exhibit a destabilized conformation, among them G130V [17], G137V [18], W173G [19] and L198R [12].

**Figure 1.**
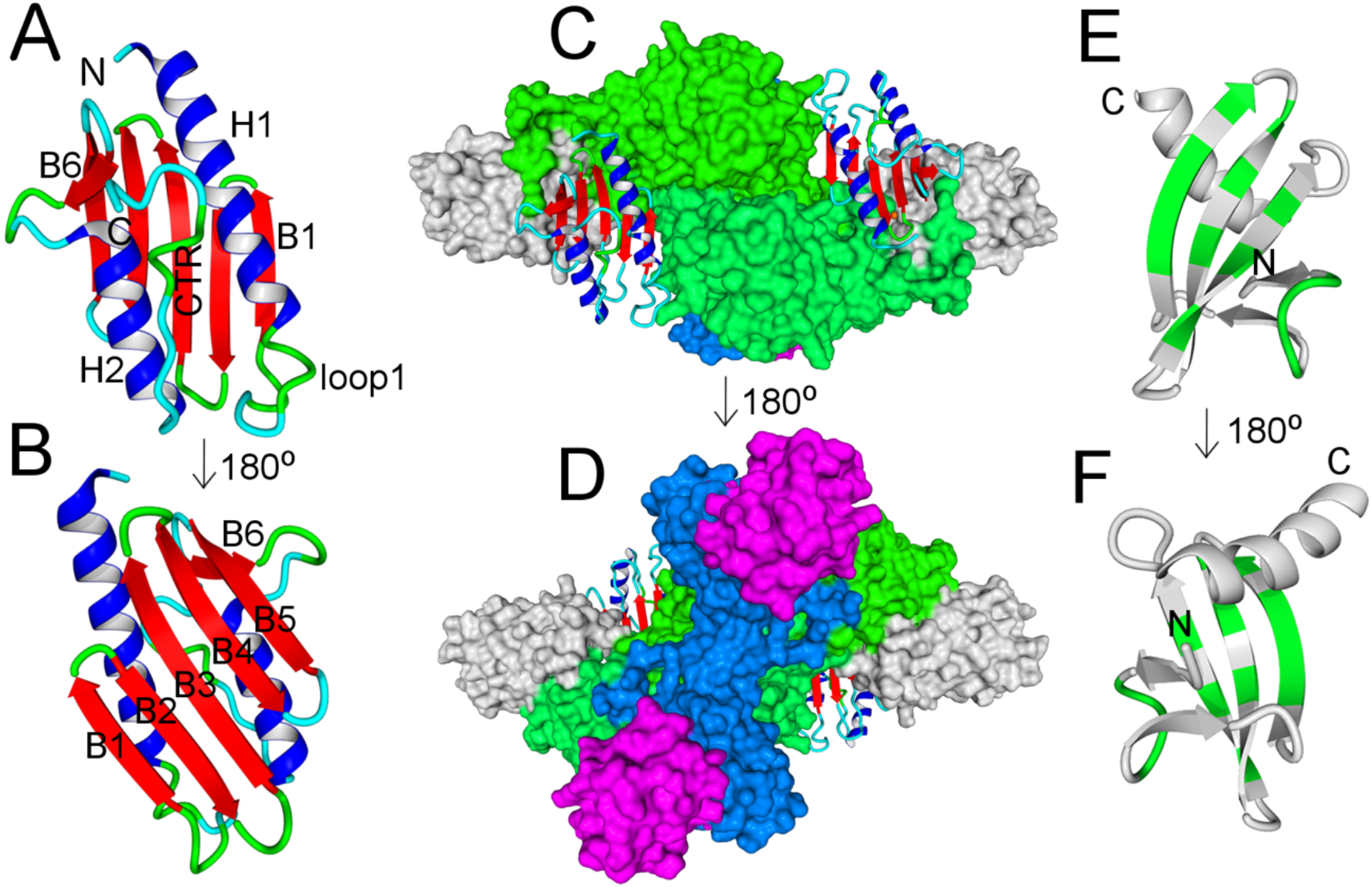
Proteins Involved in this Study. **A** and **B** are two different views of frataxin (PDB ID: 1EKG) colored by secondary structure elements. **C** and **D** correspond to the supercomplex (PDB ID: 6NZU); NFS1, ACP, ISD11 and ISCU are in green, magenta, blue and gray, in surface representation, whereas frataxin structure is in a ribbon colored by the secondary structure a. **E** and **F** are two different views of affitin (ribbon representation), whereas green residues are those residues that were randomized (positions 8, 9, 10, 11, 21, 22, 23, 24, 26, 29, 31, 33, 40, 42 and 44).

The supercomplex for iron–sulfur cluster assembly is formed by several proteins (**Figures 1C and D**) [20–22]. Two cysteine desulfurase NFS1 subunits form a dimer that is stabilized by two ACP-ISD11 heterodimers. ACP is the mitochondrial acyl carrier protein, and ISD11 is a very small protein that is only present in eukaryotic cells [23]. ISD11 subunits directly interact with the cysteine desulfurase NFS1. ACP confers stability and solubility to ISD11 [24]; in fact, the core of each ISD11 subunit is stabilized by an acyl chain that is bond via a phosphopantetheine moiety to the ACP. The structure of the isolated ACP-ISD11 is virtually identical to that found in the context of the supercomplex [25], however, in the supercomplex, ISD11 subunits self–interact and the acyl chains establish some interactions even with the Cys desulfurase NFS1. A key player in the supercomplex is ISCU, known as the scaffolding protein. One ISCU subunit interacts with each one of the NFS1 subunits. The iron–sulfur clusters are assembled on the ISCU assembly site formed by residues Cys69, Asp71, Cys95 and His137 on the surface of the protein; it is noteworthy that ISCU is able to bind 1 equivalent of iron (II) by means of these key residues [26]. Furthermore, the ferredoxin system (FdxR/FDX2) provides the electrons necessary to reduce the persulfide groups, to form the [2Fe-2S] cluster [26, 27]. Finally, the supercomplex contains two frataxin subunits. Each one of them interacts with both NFS1 subunits and with one of the ISCU subunits [22].

Thus, the architecture of the supercomplex for iron–sulfur cluster assembly is rather intricate. Moreover, the dynamics of cysteine desulfurase and the catalysis of the NFS1 enzyme are constrained by the interactions with the rest of proteins. In fact, in the absence of the ACP-ISD11 heterodimer, the enzyme is inactive [25] and monomeric [28]. On the other hand, in the presence of ACP-ID11, NFS1 gains cysteine desulfurase activity [25] and becomes dimeric [28]. Specific mutations that alter frataxin-ISCU interaction like W155R or N146K cancel frataxin–mediated activation [2, 6]. In a similar fashion, the laboratory–designed variant S157I showed highly reduced kinetic activation [29].

Considering the particular molecular architecture of the supercomplex, it is not obvious whether frataxin might simultaneously interact with other proteins and if the dynamics of these protein–protein interactions can be modulated. In fact, it is unclear whether frataxin remains in a bound state during the catalytic cycle. Based on molecular dynamics simulations performed by our laboratory, one of our working hypotheses is that wild–type frataxin forms part of the assembly site but it is also able to constrain the molecular motions of NFS1 and ISCU [19].

With the aim of generating new tools to help in understanding the biochemistry of this metabolic pathway, and ultimately, to explore new therapeutic strategies for FRDA, we have investigated the possibility of obtaining small proteins exhibiting a high affinity for frataxin. These molecules, exhibiting the capability of specifically binding to frataxin, may alter the interaction with the supercomplex and the activity of the latter. Moreover, the high affinity of the interaction between affitin and frataxin might alter the degradation of unstable frataxin variants, working as macromolecular tutors or simply modulating NFS1 activity by altering the dynamics of the enzyme. We have named this strategy “protein function stabilization by quaternary addition”.

To explore this strategy, we decided to select and study affitins (Mouratou et al., 2012) (**Figures 1E and F)** with high affinity for frataxin. We applied the ribosome display strategy, which briefly consists of three steps:

a. The generation of a high diversity synthetic library of affitins (Sac7D variants) by DNA assembly using degenerated oligonucleotides.
b. *In vitro* transcription and translation, stable ternary complexes formed by ribosome-mRNA-nascent affitin, which can interact with frataxin that is bound to a multi–well plate, are obtained.
c. A selection, whereby the complexes containing affitins with affinity for frataxin will remain bound to the plate, after a series of washings, and the mRNA is then recovered and retrotranscribed to cDNA to start a new round of affitin selection.

Several rounds (b-c) were performed, an increase of affinity and a reduction of the library diversity was expected [30]. In principle, after that, one would be able to clone the DNA fragments corresponding to Sac7D variants exhibiting a high affinity against the target.

Remarkably, the ribosome display strategy is not restricted by cell transformation and it begin with a very high diversity of sequences obtained by a PCR–based randomization step [30, 31]. This approach was useful to obtain high affinity Sac7D variants against the thermophilic CelD enzyme from *Clostridium thermocellum* and lysozyme from a hen egg [32], also against human immunoglobulin G [33] and against secretin PulD, a component of the pullulanase type II secretion system of *Klebsiella oxytoca* [34], among other protein targets. Recently, affitins have also been found of interest for therapeutic and biotechnological purposes [35–37].

In this study, we have focused on one of the affitins that we were able to select after five rounds of ribosome display using human mature frataxin as the target. We have studied affitin–frataxin interaction and discussed some applications of these new molecular tools concerning the modulation of supercomplex activity by using these small specific protein tutors that are able to bind frataxin variants.

## MATERIALS AND METHODS

### Library Assembly, Affitin Selection and Purification

Ribosome display for the selection of Sac7d variants was carried out as outlined by Mouratou and coworkers (Mouratou et al., 2012). Primers used for the DNA library assembly were purified by HPLC. The mRNA was prepared *in vitro* by using the HiScribeTM T7 high–yield RNA synthesis kit (New England Biolabs) and purified using an RNeasy MinElute Cleanup kit (QIAGEN). The RNAse inhibitor Ribolock (Thermo Fisher) was added, whereas the RNA working zone was decontaminated using RNAse Away (Thermo Fisher) to avoid DNAse and RNAse presence.

*In vitro* protein translation was carried out (in absence of the release factors) using the PURExpress ΔRF123 kit (New England Biolabs). This guaranteed the stabilization of the ternary complexes mRNA-affitin-Ribosome. After each cycle of affitin selection, the recovered mRNA was retrotranscribed using SuperScript III reverse transcriptase (Thermo Fisher); the RNA was digested using DNAse–free RNAse 1 (Thermo Fisher) and the cDNA was amplified by PCR using Vent DNA polymerase (New England Biolabs). After DNA purification using the QIAEX II kit (QIAGEN), the DNA was either used to prepare a new library for the next selection cycle or cloned (see below).

For each round of selection, frataxin (100μL of 1-5 μg/mL) was directly bound to the MaxiSorp microtiter plates (NUNC), and BSA (bovine serum album, SIGMA) was used to block the remaining sites (overnight incubation at 4 °C). The elution buffer was 50 mM Tris-acetate pH 7.4 at 4 °C, 150 mM NaCl, 20 mM EDTA (sterilized by 0.22 μm filtration). Yeast RNA (Roche) was included as the vehicle for RNA recovery.

To increase the chance of finding affitins with a high affinity to frataxin, we performed prepanning steps against BSA. Additionally, we reduced the frataxin concentration on the plate surface and the number of PCR cycles after the retro transcription step.

For the last two selection cycles, the cDNA obtained was cloned in plasmid pFP1001 (derived from pQE30, QIAGEN) using BamHI and HindIII restriction enzymes and T4 DNA Ligase (Thermo Fisher). *E. coli* DH5 alpha F’IQ competent cells were transformed and affitin clones were isolated. For clone selection, plasmid amplification and protein expression, 100 μg mL^−1^ ampicillin and 25 μg mL^−1^ kanamycin were added to the culture medium. In particular, for protein expression, 2YT 0.1% glucose was used, and induction was carried out when DO_600nm_ reached 0.75 by the addition of 0.5 mM IPTG by overnight incubation at 30 °C under agitation at 200 rpm. Affitins were N-terminal tagged with the RGS-His6 sequence for detection and purification. The affitin detection was carried out using the RGS-His6-HRP conjugate (QIAGEN).

After selection, the isolated clones were screened by ELISA for affitin–frataxin interaction. For this analysis, protein induction was performed on a small scale (2mL) and cells were disrupted using BugBuster protein extraction reagent (Novagen).

The selected clones were evaluated both for frataxin interaction and for BSA as a control, because the latter was also added to the selection to block the plates. Even though a prepanning step had been performed, it was expected that some clones of affitins exhibiting high affinity to BSA might be selected. DNA corresponding to the affitin clones was sequenced by the Macrogen facility.

For frataxin–affitin interaction detection by ELISA, *E. coli* BugBuster extracts, a soluble fraction after sonication and subsequent centrifugation, or purified proteins were all methods used to evaluate the interaction between affitins and human frataxin. As we have mentioned above, Maxisorp plates (NUNC) coated with BSA were used to establish the specificity of affitins for frataxin. The detection of binding was carried out by the RGS His6 HRP conjugate (QIAGEN) that detects the RGS His6 motif present in the affitins N-terminal. Typically, a dilution of the antibody of 1:4000 in TBS-Tween 0,1% was used for ELISA detection. The horseradish–peroxidase substrate was 3,3’,5,5’-tetramethylbenzidine (TMB, Thermo Fisher). The absorbance at 450 nm was read after the addition of 4N sulfuric acid.

For affitin variant purification, the pellet corresponding to 1-2L of affitin–induced cell culture was resuspended in 15mL of 50mM Tris-HCl, 200 mM NaCl, pH 7.5 buffer. Cells were disrupted by sonication, and the recombinant protein was purified from the soluble fraction (separation after 30 min 10,000 rpm) using a Ni^2+^-NTA-agarose column (GE Healthcare), equilibrated in 50 mM Tris-HCl, 300mM NaCl, pH 7.5 buffer. The affitins were eluted by means of increasing concentrations of imidazole prepared in the same buffer. Fractions containing the recombinant protein, as judged by the SDS-PAGE analyzed, were pooled and dialyzed against a buffer without imidazole and containing 1-5 mM DTT, if necessary. Identity was verified by mass spectrometry analysis (ESI-MS).

### Frataxin Production and Purification

Wild–type frataxin (residues 90-210) and mutants were expressed and purified as previously described for the wild–type variant [12]. Briefly, frataxin 90-210 was subcloned into a pET9b plasmid and cells *E. coli* BL21 (DE3) were transformed. Bacteria cultures 2–3 L (LB medium, pH 7.2) were grown at 280 rpm and 37 °C. Protein expression was induced at DO= 1.0 by the addition of 1% (*w/v*) lactose (final concentration) and induction was carried out for 4 hours. Bacteria were centrifuged at 6,000 rpm and the pellet was stored at –20 **°**C until cell disruption by sonication in a 20 mM Tris-HCl buffer, 1mM EDTA, pH 7.0. After centrifugation at 10,000 rpm (30 min), frataxin (~100%) was found in the soluble fraction. For protein purification the soluble fraction was carefully loaded onto ion exchange chromatography (DEAE DE52 matrix) and eluted with a 300 mL linear gradient (from 0.0 to 1.0 M NaCl) prepared in a 20 mM Tris-HCl buffer, 1 mM EDTA, pH 7.0. Afterwards, fractions (identified by SDS-PAGE) with frataxin were loaded onto a preparative Sephadex G–100 column (SEC, 93 cm62.7 cm), previously equilibrated with a 20 mM Tris-HCl buffer, 100 mM NaCl, 1 mM EDTA, pH 7.0. This protocol yielded approximately >98% pure frataxin as confirmed by SDS-PAGE and mass spectrometry (ESI-MS); the theoretical molecular mass value of frataxin was 13,605.1 Da. Frataxin concentration was determined spectroscopically using an extinction coefficient ε_280nm_ = 26,930 M^−1^ cm^−1^ (1 mg/mL protein solution represents Abs_280nm_ = 2.00).

Identity was verified by DNA sequencing using the Macrogen facility and by mass spectrometry analysis (ESI-MS). The latter was performed in the National Laboratory of Research and Services in Peptides and Proteins using an LCQ DUO ESI ion trap (Thermo Finnigan) or QExactive Orbitrap (Thermo Scientific) spectrometers.

### Expression of Human Cysteine Desulfurase NFS1, Human Acyl Carrier Protein, ISD11 and ISCU2

The DNA sequences corresponding to the human mature form of the human cysteine desulfurase NFS1 enzyme (NFS1Δ55), the human mitochondrial acyl carrier protein (ACP, the mature form) and the ISD11 were optimized for protein expression in *E. coli* BL21 (DE3) by BIO BASIC Inc (Markham ON, Canada). NFS1Δ55 and ISD11 were cloned in a pETduet plasmid, whereas ACP was cloned in a pACYCDuet-1 for co-expression. The NFS1 amino acid sequence included the RSGHHHHHH tag in the N-terminal for purification and antibody recognition. Co-expression (1mM IPTG induction at OD_600nm_=0.8-1.0) was carried out overnight at 20 °C and 250 rpm.

The purification of the complex (NFS1/ACP-ISD11)_2_ was performed from the soluble fraction using a Ni^2+^-NTA-agarose column (first step). The complex was eluted with 20 mM Tris-HCl, 300 mM NaCl, 500 mM imidazole, pH 8.0. The fractions were analyzed by SDS-PAGE; fractions showing a high concentration of the complex were pooled and frozen at −20 °C. Before using, the complex was thawed in ice and centrifuged to remove possible protein aggregates; nevertheless, the co-expression of the three proteins yielded a very stable complex. On the other hand, purification of NFS1 alone (in the absence of ISD11 and ACP) resulted in a more unstable protein sample, with a higher tendency to aggregate, as previously shown [3, 5]. This is why we ultimately preferred the co-expression strategy rather than reconstitution by mixing NFS1 and ACP-ISD11.

The plasmid, pET-22b, encoding the human ISCU2, i.e., the scaffolding protein (Uniprot ID: Q9H1K1-1, C-terminal His-tagged) of the supercomplex, was generated by Explora Biotech (Rome Italy). *E. coli* BL21(DE3) cells were transformed. Protein expression was carried out in a LB medium containing 50 μg/mL ampicillin by adding 1% (w/v) lactose at OD_600 nm_ = 0.8. After 4 h of culture at 28 °C, cells were harvested by centrifugation and stored at −20 °C. ISCU purification was performed in a similar way to the one described in the previous paragraph for cysteine desulfurase NFS1, but the pH value was set at 7.5.

### Protein Characterization by Circular Dichroism

Circular dichroism (CD) measurements were carried out in a Jasco J-815 spectropolarimeter. Far– and near–UV CD spectra were collected using 0.1 and 1.0 path length cells, respectively, thermostatized at 20 °C. Protein concentrations were between 5 and 50 μM and the buffer was 20 mM Tris-HCl, 100 mM NaCl, pH 8.0. Data was acquired at a scan speed of 20 nm min^−1^, with a 1-nm bandwidth and a 0.1-nm data-pitch. Five scans for each sample were averaged, and the blank spectrum was subtracted.

### Affitin Characterization by SEC-FPLC

The oligomerization state and molecular mass determinations were carried out by Multiple-Angle Laser Light-Scattering (MALLS) using a miniDawn (Wyatt Technology) coupled to a size–exclusion column (SEC, Superose-12, GE Healthcare), at room temperature and at a 0.3-0.4 mL min^−1^ flow rate. Data analysis was carried out with Astra 6.0 software (Wyatt Technology). Protein concentration was 100 μM and the buffer was 20 mM Tris-HCl and 150 mM NaCl, at pH 7.5. A concentration of 1mM DTT was added when necessary.

### NFS1 Cysteine Desulfurase Activity

Enzymatic desulfurization of cysteine to alanine and sulfide by the (NFS1/ACP-ISD11/ISCU/FXN)_2_ supercomplex was determined by the methylene blue method. Concentrations of proteins, substrate and the reducing agent (DTT) were set according to a previous paper by Tsai and Barondeau [5].

Reactions contained 1.0 μM NFS1, 1.0 μM ACP-ISD11, 3.0 μM ISCU and 3.0 μM FXN, and samples were supplemented with 10 μM PLP, 2 mM DTT, and 1.0 μM FeSO_4_ (final concentrations). The reaction buffer was 50 mM Tris-HCl and 200 mM NaCl, pH 8.0. Reactions were started by the addition of 1 mM cysteine (K_m_ for cys is 10-20μM). Samples were incubated at room temperature for 30 min. H_2_S production was stopped by the addition of 50 μL of 20 mM N,N-dimethyl p-phenylenediamine in 7.2 M HCl and 50 μL of 30 mM FeCl_3_ (prepared in 1.2 M HCl). Under these conditions, the production of methylene blue took 20 min. After that, samples were centrifuged for 5 min at 12000*g*, and the supernatant was separated. Absorbance at 670 nm was measured.

### Affitin-Frataxin Interaction Perturbation Experiments Followed by TEXAS-RED Fluorescence

The fluorescence assay consisted in direct immobilization of affitin 224 (100 μl of 10 ng/μl in TBS buffer, 4*C ON incubation) in a black–flatted multi–well plate (NUNC). The plate was then blocked with a solution of 1% BSA in a TBS buffer for 2 h at room temperature. After several washes with a TBS-0.05% Tween buffer, a Texas Red–labeled S202C FXN variant was incubated under different conditions for 1 hour at room temperature. Several washes were performed with a TBS-0.1% Tween buffer and then fluorescence reading was performed in a Tecan Instrument (excitation and emission wavelengths were 590 and 630 nm, respectively) in the presence of a 100 μl of TBS buffer. The S202C FXN variant was previously labeled with Texas Red maleimide dye (Thermo Fisher) as previously described by Das et al. [38] and manufacturer recommendations.

### Surface Plasmon Resonance

Affinity measurements were performed using BioNavis VASA 210 equipment (BioNavis Ltd, Tampere, Finland). Carboxymethylated dextran sensors were purchased from Sofchip (Sofchip, Florida, United States) and used for frataxin immobilization. Briefly, immobilization was performed *in situ* in the instrument. The surface was washed with 2M NaCl and 50mM NaOH, and with Milli Q water. The surface was then activated by the injection of 100 mM EDC (1-ethyl-3-(3-dimethylaminopropyl)carbodiimide) and 50 mM NHS (N-hydroxysuccinimide) using a 10 mM acetate buffer pH 4.1 for 7 minutes. The solution was freshly prepared before use. Then a solution of 7 μM frataxin in a sodium acetate buffer 10 mM pH=4,0 was injected for 7 minutes in channel 1 at a flow rate of 20 μL min^−1^. The reference surface was created in channel 2 by injecting a buffer solution. Subsequently, the unbound sites were blocked using 1.0 M Ethylene diamine for 7 min at a flow rate of 30μL min^−1^.

The experiments were performed in a 20 mM Tris-HCl, 150 mM NaCl, 1 mM DTT, pH 7.5 measurement buffer at 21 °C and a flow rate of 30 μL min^−1^. The buffers were previously degassed. The sample injections were performed for 5 min for the association phase and 10 minutes for the dissociation one. Serial dilutions of affitins were analyzed, as is required for the kinetic analysis of molecular interactions. The regeneration of the surface, using a solution of 50 mM NaOH, was performed between each sample injection. Measurements were performed in triplicate.

Sensogram analysis were performed using DataViewer Software from BioNavis Ltd. Each sample was referenced using a blank channel (channel 2). Kinetic analysis was performed using TraceDrawer 1.3 for BioNavis Ltd. A 1–to–1 interaction model was fitted to the curves obtained.

### Affitin–Frataxin Interaction Studied by ^1^H ^15^N NMR

Samples for NMR experiments contained 0,1 mM ^15^N labelled hFXN (90-210) and 0,17 mM unlabeled Affi_224 in a buffer containing 20mM Tris-HCl, 100mMA NaCl, 0,5mM TCEP, pH 7.0 and 5% D_2_O. NMR experiments were performed at 22 °C in a Bruker 600 MHz Avance III spectrometer (Bruker Instruments, Inc., Bellerica, MA, USA). ^1^H-^15^N heteronuclear single quantum coherence (HSQC) spectra of ^15^N labelled FXN (variant including residues 90-210) were obtained and compared with previous ones [12, 14, 15]. The NMR data were processed with NMRPipe [39] and analyzed using NMRViewJ [40]. Chemical shift perturbations (CSP, [41]) were calculated according to Equation 1:

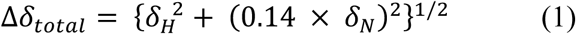

where the shift changes in ^1^H and ^15^N are *δ*_*H*_^2^and *δ*_*N*_, respectively.

### Cell Culture and Protein Extraction: Affitin 224 Expression in HEK-293T Cells

Affitin transfection and expression was carried out in HEK-293T cell line (kindly provided by Dr. Itatí Ibanez, INQUIMAE, UBA). HEK-293T cells were grown in a high glucose (4.5 g L-1 glucose) Dulbecco’s modified Eagle’s medium (DMEM, Thermo Fisher Scientific) supplemented with 10% fetal bovine serum (FBS, Natocor), penicillin/streptomycin (100 units mL^−1^ and 100 μg mL^−1^ respectively, Thermo Fisher Scientific) and 110 mg L-1 of sodium pyruvate (Thermo Fisher Scientific) in a 37 °C humidified incubator containing 5% CO_2_. Cells were plated (2 ×10^6^ cells per 100 mm plate) and grown for 24 h before transfection with Polyethylenimine (PEI, Sigma) according to the manufacturer’s instructions. Cells were grown for 48 h before harvesting (by using RIPA buffer) to check affitin expression. Total protein concentration in the samples was quantified using a Bradford reagent (Biorad).

Successful transfection and affitin expression were checked by Western blotting using a RGS-His6 HRP conjugate (Qiagen) to detect Affi_224 in a HEK-293T cell lysate. Previously, cell viability was estimated by the dye exclusion test proposed by Strober W [42], counting cells in a Neubauer Chamber that include (dead cells) or exclude (live cells) Trypan blue dye (Sigma).

### Affitin 224-Frataxin Co-Immunoprecipitation Assay

250 μg of protein from HEK-293T cell lysates were incubated overnight with 2 μg of a 6x-His tag antibody (MA 1-135, Thermo Fisher) in the presence of a low–salt RIPA buffer (20mM Tris-HCl, 150mM NaCl, 0,1% between 20, 2 mM EDTA, pH= 7.4) and complete protease inhibitor (Thermo Fisher). The binding to protein G magnetic beads (Biorad) was performed for 1 hour at 4 ºC (with previous washes with a RIPA buffer). Three washes were performed after binding and then eluted with 25 μl of a 1X SDS-PAGE sample buffer containing 8% Beta-Mercaptoethanol. Mouse IgG Isotype control was performed using anti HPRT (Santa Cruz Biotechnology). The IP assays using the human FXN 81-210 variant and pure Affi_224 were performed in similar conditions.

### SDS-PAGE and Western Blotting Analysis

Protein samples (either purified or complete lysates) were boiled in a sample buffer (4% SDS, 20% glycerol, 120 mM Tris, pH 6.8, 0.002% bromophenol blue, 200 mM 2-mercaptoethanol) and subjected to a homemade 16% SDS-PAGE. Electrophoresis was carried out at room temperature for 20 min at 90V and 1.5 hours at 150V. Proteins were stained with Coomassie Brilliant Blue G-250. When a Western blot analysis was performed, proteins were transferred to a PVDF membrane (0.2 μm, Thermo Fisher) during 1 h at 100 V. Membranes were blocked for 1 hour at room temperature either with a commercial blocking buffer (QIAGEN, anti RGS HIS6 HRP conjugate kit) or with 5% milk in 0.05% tween TBS buffer. Blocked membranes were incubated overnight at 4 °C with either an anti human frataxin monoclonal antibody (abcam, ab 110328), an anti HPRT monoclonal antibody (SC-376938), an anti GAPDH monoclonal antibody (SC-47724) or an anti RGS HIS6 HRP (QIAGEN). When necessary, HRP-conjugated anti-mouse was incubated for 1 h at room temperature and visualized by enhanced chemiluminescence (Clarity Substrate, Biorad) using Amersham Imager 680.

## RESULTS AND DISCUSSION

### Selection, Expression and Conformational Characterization of Frataxin Binders

Using the Ribosome Display strategy proposed by Mouratou and coworkers [30], a total of fifteen positions of the affitin were randomized (positions 8, 9, 10, 11, 21, 22, 23, 24, 26, 29, 31, 33, 40, 42 and 44) and more than 400 clones were isolated after completion of the fifth cycle of selection. These clones were analyzed for affitin–frataxin interaction by ELISA. Some clones showed to be specific for frataxin. On the other hand, some clones exhibited interaction only against BSA that was used to block the surface of the selection plates.

Soluble fraction of *E. coli* crude extracts expressing five different affitins exhibited high frataxin/BSA signal ratios (**Fig. 2A**). This result suggested that the interaction is specific, and the presence of other proteins does not interfere with the interaction. For comparison, Affi_12 is an example of an unspecific affitin.

**Figure 2.**
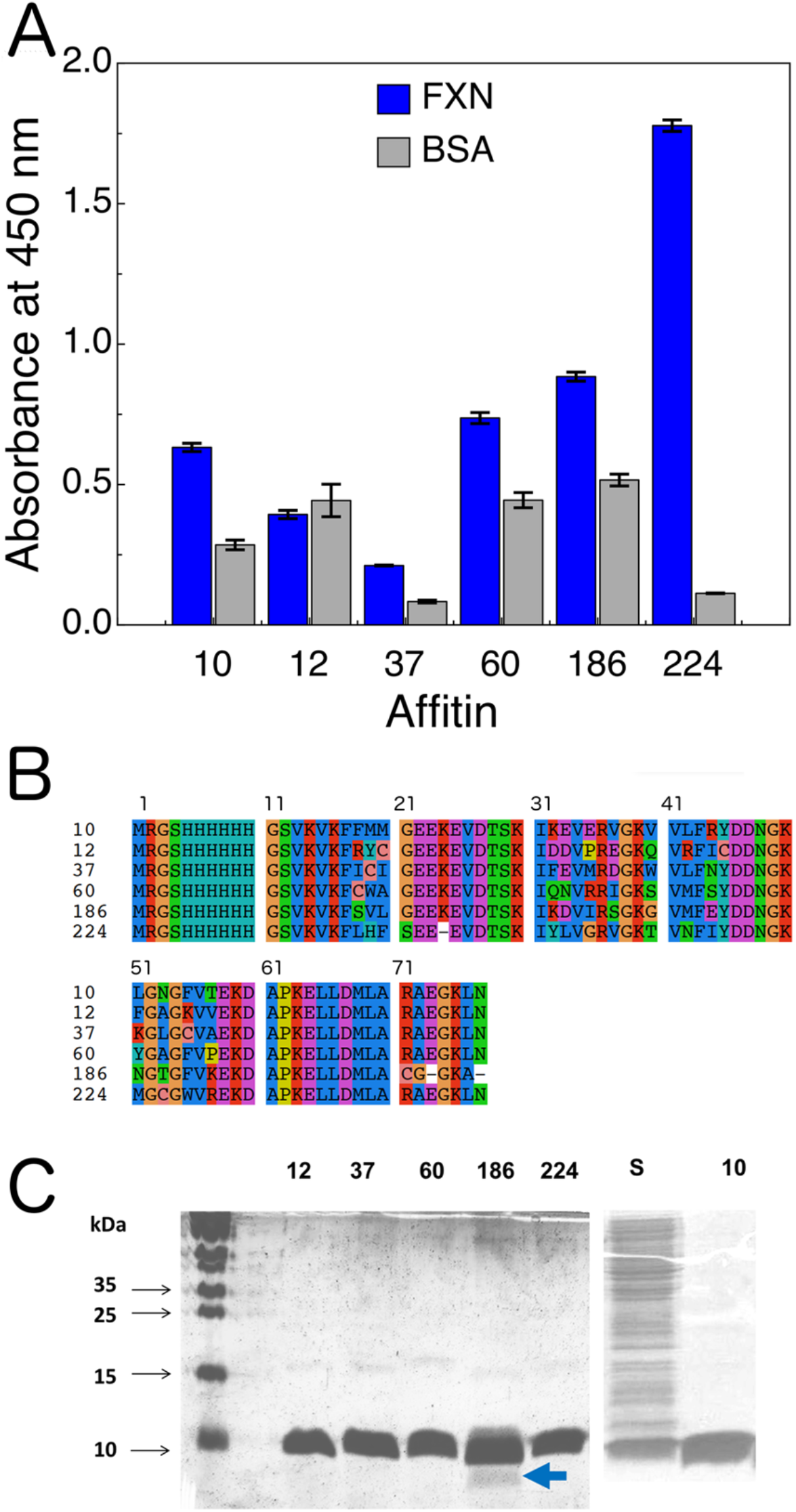
The Selected Affitins. **(A) ELISA Screening of Positive Clones of Round 4 and 5.** The positive clones from these rounds of selection were assayed together in the same assay. The induced clones were subjected to lysis and the supernatant was diluted with TBS. Affitin dilutions were incubated for 1h at room temperature in wells previously coated with frataxin or directly coated with BSA. After that, the plate was washed with TBS and then incubated with an anti RGSHHHH peroxidase–conjugated antibody (dilution 1:4000) to detect affitin–target interaction. TMB substrate was added. Absorbance was read at 450 nm. (B) Amino acid sequences of the selected affitins, deduced from the DNA sequence. (C) Analysis of the selected affitins by 16% SDS-PAGE after expression and purification. The blue arrow indicates partial degradation of affitin 186.

In **Fig. 2B** we show the amino acid sequences corresponding to these clones. Even though Affi_60, Affi_186 and Affi_224 showed a high expression under these experimental conditions, Affi_37 showed a significantly lower protein expression level. This fact was confirmed when affitin Affi_37 was expressed for protein purification. Affitins were purified and in all cases, purity was >95% as judged by the SDS-PAGE (**Fig. 2C**) and mass spectrometry analysis. Purified affitins interacted with frataxin directly attached to the plates.

It is worthy of note that 5/6 of the selected affitins contain at least one cysteine residue in their amino acid sequences. The analysis by mass–spectrometry (ESI-MS, **Table 1**) showed that at least Affi_186 and Affi_224 formed intermolecular disulfide bonds. High concentrations of DTT followed by chemical modification with iodoacetamide showed peaks shifted by +57Da, compatible with the addition of the -CH_2_-CO-NH_2_ moiety.

**Table 1.** Molecular Properties of Affitins.

| **Affitin** | **Molecular Mass<sup>1</sup>**<br>**(Da)** | **Experimental**<br>**Molecular**<br>**Mass (Da)** | **Isoelectric**<br>**Point**<br>**1** | **Cysteine**<br>**content<sup>1</sup>** | **Extinction**<br>**coefficient**<br>**(M<sup>-1</sup>cm<sup>0</sup><sup>-1</sup>)** |
| --- | --- | --- | --- | --- | --- |
| 10 | 8790.07 | 8789.3 | 8.07 | 0 | 1490 |
| 12 | 8712.86 | 8582.1<sup>2</sup><br>(-130.76)<br>8598.2<sup>2</sup><br>(-130.76<br>+16.1) | 7.85 | 2 | 1490 |
| 37 | 8732.01 | 8730.2<sup>2</sup> | 7.01 | 2 | 6690 (7115) |
| 60 | 8706.86 | 8708<sup>2</sup> | 8.79 | 1 | 8480 |
| 186 | 8249.36 | 8305.41<sup>3</sup><br>(+56.04)<br>16497<sup>4</sup> | 7.92 | 1 | 1490 |
| 224 | 8687.90 | 8744.5<sup>3</sup><br>(+56.6)<br>17374<sup>4</sup> | 7.96 | 1 | 8480 |

<sup>1</sup> Calculated from the DNA sequence.
<sup>2</sup> Determined by ESI-MS after reduction with DTT.
<sup>3</sup> Determined by ESI-MS after reduction with DTT and iodoacetamide modification (+57Da).
<sup>4</sup> Determined by ESI-MS without DTT.

In the case of Affi_12, peaks shifted by ~131 Da (130.76 Da lower), a mass compatible with the absence of Met1. For this affitin, a peak shifted by 16 Da (+16.1) was also present, compatible with the oxidation of the Met 68 residue.

The SEC-FPLC analysis indicated that the Affi_10, Affi_60, Affi_186 and Affi_224 behave as dimers in equilibrium with monomers, in solution (**Fig. 3**). This occurred even in the presence of 1.0 mM DTT. It is worthy of note that Affi_10 does not include Cys residues, showing that the selected affitins have a marked tendency to dimerize independently of the oxidation state of the proteins.

**Figure 3.**
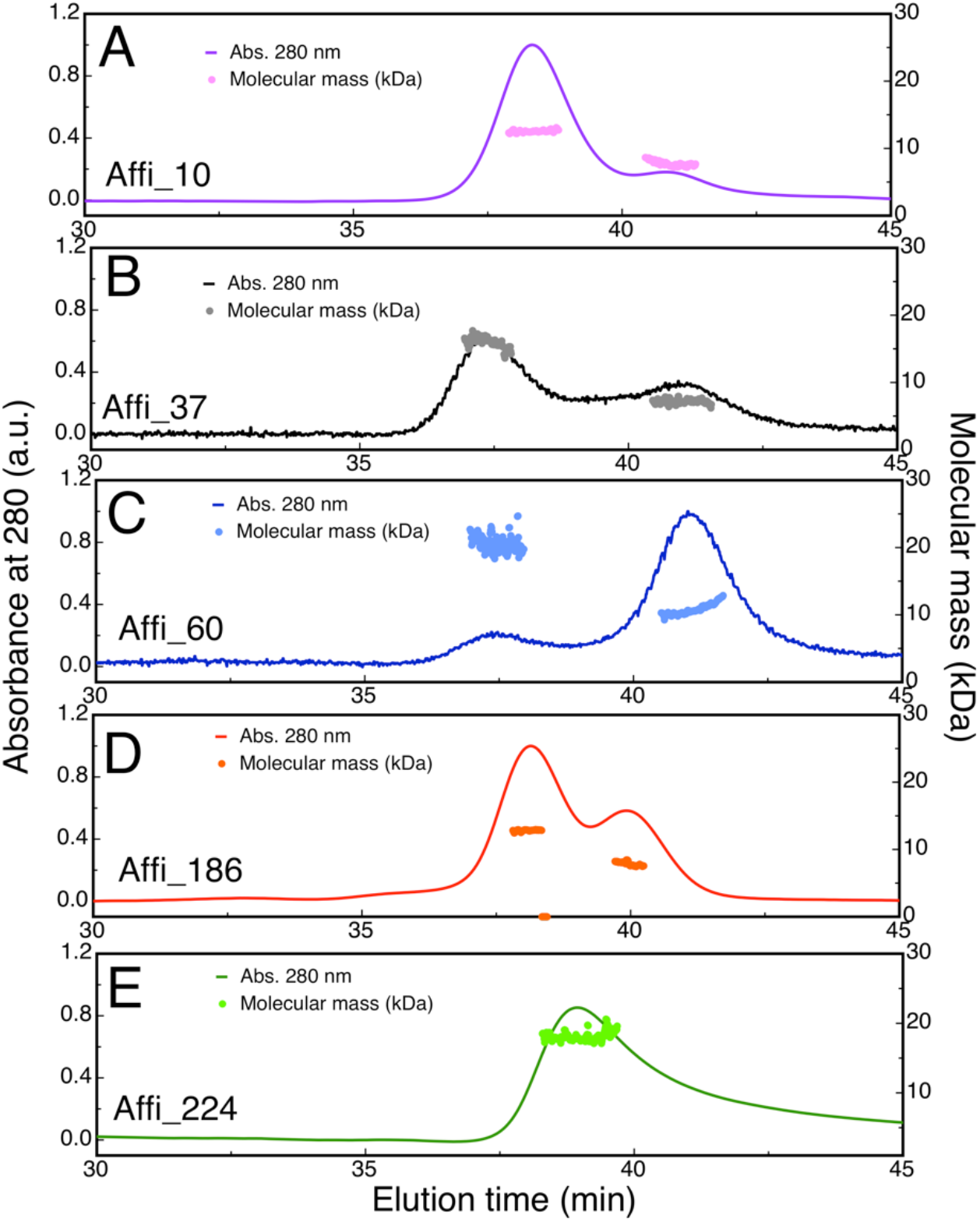
Analysis of the Affitins by SEC and MALLS. Protein concentration was 100 μM and the buffer was 20 mM Tris-HCl and 150 mM NaCl, 1mM DTT, at pH 7.5. MALLS and UV detectors were coupled to a Superose-12, (GE Healthcare). The experiment was carried out at room temperature.

Molecular mass of affitins in solution determined by multi–angle light scattering was also indicative of the presence of at least two species (dimer and monomer). In the case of Affi_186, a lower molecular way than expected was obtained, probably as a consequence of the uncertainty of the extinction coefficient; it is worthy of note that the observed mass (ESI-MS) is consistent with what was expected for the full–length Affi_186.

On the other hand, Affi_224 exhibited a different hydrodynamic behavior. Its SEC profile suggested a faster equilibrium between dimeric and monomeric species than the values observed for the other affitins.

We tried to block the Cys residues under native conditions by using alkylating agents to cancel the possibility of disulfide bond formation and to evaluate monomer–dimer equilibrium. The results indicated that, at least for the case of affitin Affi_224, the modification of the Cys residue with iodoacetamide highly increased the tendency of the protein to aggregate; after protein modification, ~75% of the protein precipitated. This fact was also evident when Cys residue was modified with Texas red dye, precluding the evaluation binding monitored by fluorescence of Texas red–labeled Affi_224.

To investigate the location of the Cys residue/es in the affitin structures, we prepared molecular models using the *Swiss model* (Waterhouse et al. 2018). Models showed that for Affi_60, Affi_224 and Affi_186, the cysteine residue would be located on the surface of the protein, whereas for Affi_37 the two cysteine residues may be only partially exposed to the solvent (**Fig. S1**). Thus, the formation of disulfide bonds in the context of structured affitin dimers might be possible.

Moreover, we studied the degree of structure of the affitins by circular dichroism (CD) (**Fig. 4**). With the exception of Affi_186, the near-UV CD spectra indicated that the aromatic residues of these proteins are located in asymmetric environments as a product of the native structure (**Fig. 4A**). Similarly, the far-UV CD spectra profiles of Affi_37, 60 and 224 were compatible with structured proteins (**Fig. 4B**).

**Figure 4.**
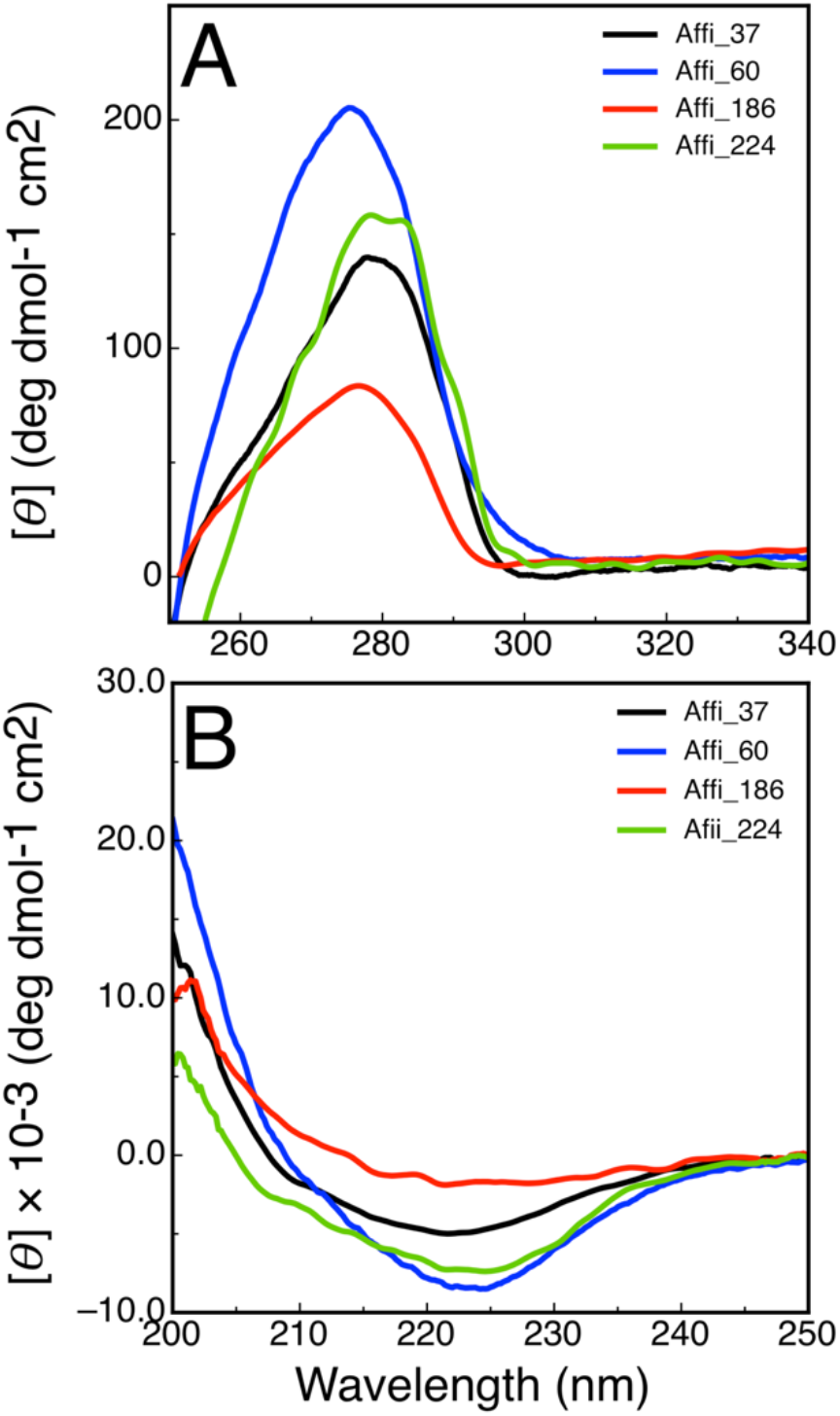
Affitin Characterization by Circular Dichroism. Affi_37, Affi_60 Affi_186, Affi_224 spectra were acquired at 25 *μ*M in buffer was 20 mM Tris-HCl, 150mM NaCl and pH 7.0.

Remarkably, Affi_186 showed diminished near and far-UV CD bands (**Figs. 4 A and B**), suggesting that a significant fraction of this protein might be unfolded or partially folded, even in the absence of denaturant and at 25 °C and pH 7-7.4. On the other hand, a difference in the extinction coefficient did not justify the large difference observed in the CD spectra of Affi_186 compared to the rest of the affitins.

### Characterization of the Affitin-Frataxin Interaction

We evaluated interaction between affitins and frataxin by surface plasmon resonance (SPR). Frataxin was covalently bound to the chip surface using the amine (N-terminus and ε-amino groups of lysine residues) coupling procedure. Under these conditions, in principle, there should be no preferential orientations of the protein. As the affitins have a significant tendency to form dimers and to establish intermolecular disulfide bonds, we included 1.0 mM DTT in the running buffer. SPR sensogram profiles were analyzed and the kinetic coefficients (*k*_on_ and *k*_off_) and equilibrium constant (*K*_D_) were determined (**Table 2, Fig. 5**). Two affitins (Affi_37 and Affi_224) exhibited a *K*_D_ in the nanomolar range, whereas the other two affitins, Affi_60 and Affi_186, showed *K*_D_ values in the sub-micromolar range.

**Table 2.** Frataxin-Affitin Binding Features Evaluated by Surface Plasmon Resonance.

| Affitin | $k_{on}$<br>($M^{-1} S^{-1}$) | $k_{off}^1$<br>($S^{-1}$) | $K_D$<br>(nM) |
| --- | --- | --- | --- |
| 37 | 18600±2 | 0.000624 | 33.5±0.1 |
| 60 | 7400±8 | 0.000900 | 129.0±0.2 |
| 186 | 2130±5 | 0.000920 | 432.0±1.2 |
| 224 | 22700±1 | 0.000691 | 34.0±0.1 |

<sup>1</sup> In all cases the standard deviation is lower than $2 \times 10^{-7} S^{-1}$ .

**Figure 5.**
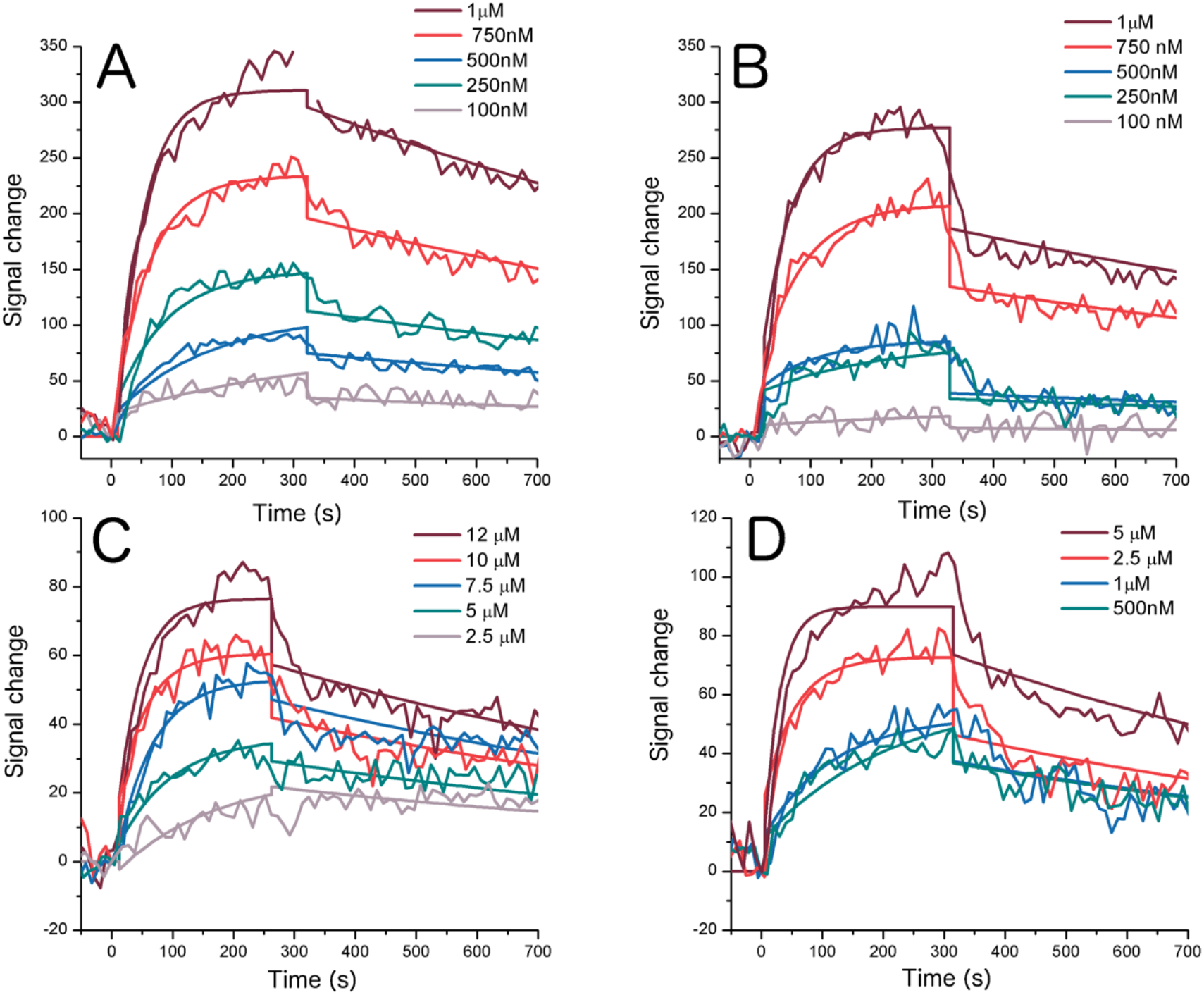
Interaction between FXN and Affitins Monitored by SPR (Surface Plasmon Resonance). (A) Affi_224, (B) Affi_37, (C) Affi_186, (D) Affi_60. Buffer was 20mM Tris-HCl, 150mM NaCl, pH 7.5, in the presence of 1mM DTT. FXN was bound to the SPR chip, whereas the affitin concentrations assayed are indicated. A one-site binding model was fitted to the data in all cases.

Given that Affi_224 showed an excellent signal ratio in ELISA (frataxin vs. BSA) and the lowest *K*_D_ value, we continued working with this protein as the affitin candidate.

Affitins accumulated several mutations in their amino acid sequence, Affi_224 showed to be marginally stable in solution. Urea–induced unfolding experiment (**Fig. S2A**) analysis indicated a conformational stability of *ΔG*°_NU_= 3.0 kcal/mol, and *m*_NU_ = 1.3 kcal/mol*M and a *C*m =2.3 M; as the parental protein Sac7D is significantly more stable (*ΔG*°_NU_= 7.3 kcal/mol at 25 °C and pH 7.0, [43], some of the mutations selected have to be destabilizing. Accordingly, Affi_224 exhibited a Tm value of 49-50 °C, as judged by temperature–induced unfolding monitored by CD at 220 nm (**Fig. S2B**).

To a further study of the interaction between frataxin and Affi_224, binding was perturbed by means of changes in the pH, or NaCl, urea or detergent concentrations (**Fig. S3**). To this end, Affi_224 was bound to a multi–well plate and Texas red–labeled frataxin was added. After incubation and subsequent washing to eliminate the remaining labeled frataxin (bound to the affitin), fluorescence was evaluated.

Interaction was substantially preserved in a broad range of pH values (using Tris-HCl buffer). It was not significantly perturbed by the addition of NaCl, in the range of 100-450 mM. Moreover, Tween-20 in the range 0-0.15% did not alter interaction. On the other hand, a concentration of 2-3 M urea prevented frataxin binding, a result consistent with the alteration of interaction, affitin unfolding or both.

Affi_224 was able to interact with frataxin directly bound to the ELISA plate and also with frataxin bound to the plate *via* biotin–neutravidin interaction. For these experiments, we prepared two frataxin variants (H177C and S202C) that were biotinylated at positions 177 or 202 respectively, **Fig. S4**). The results suggested that Affi_224 was able to interact with frataxin orientated from these two different sites.

To gain information concerning which residues of frataxin were involved in the binding process to the Affi_224, we prepared ^15^N-labeled frataxin and the interaction was studied by NMR spectroscopy by analyzing the chemical shift perturbations (CSP) under ^1^H-^15^N-HSQC experiments. As expected, the majority of the cross peaks did not present CSP when affi_224 was incubated with ^15^N-frataxin; however, some cross peaks showed significant CSP (**Fig. 6, Fig. S5**). When residues that showed CSP (Asp104, Glu108, Leu113, Ala114, Asp115, Thr119, Tyr123, Asp124, Val125 and Phe127 Gly130 and Leu132) were mapped on the frataxin structure, it was possible to observe that all of these residues were located close in space and involved residues from the acidic ridge of frataxin (formed by alpha1, loop1 and beta1 elements with a high content of Glu and Asp residues, **Fig. 7D**).

**Figure 6.**
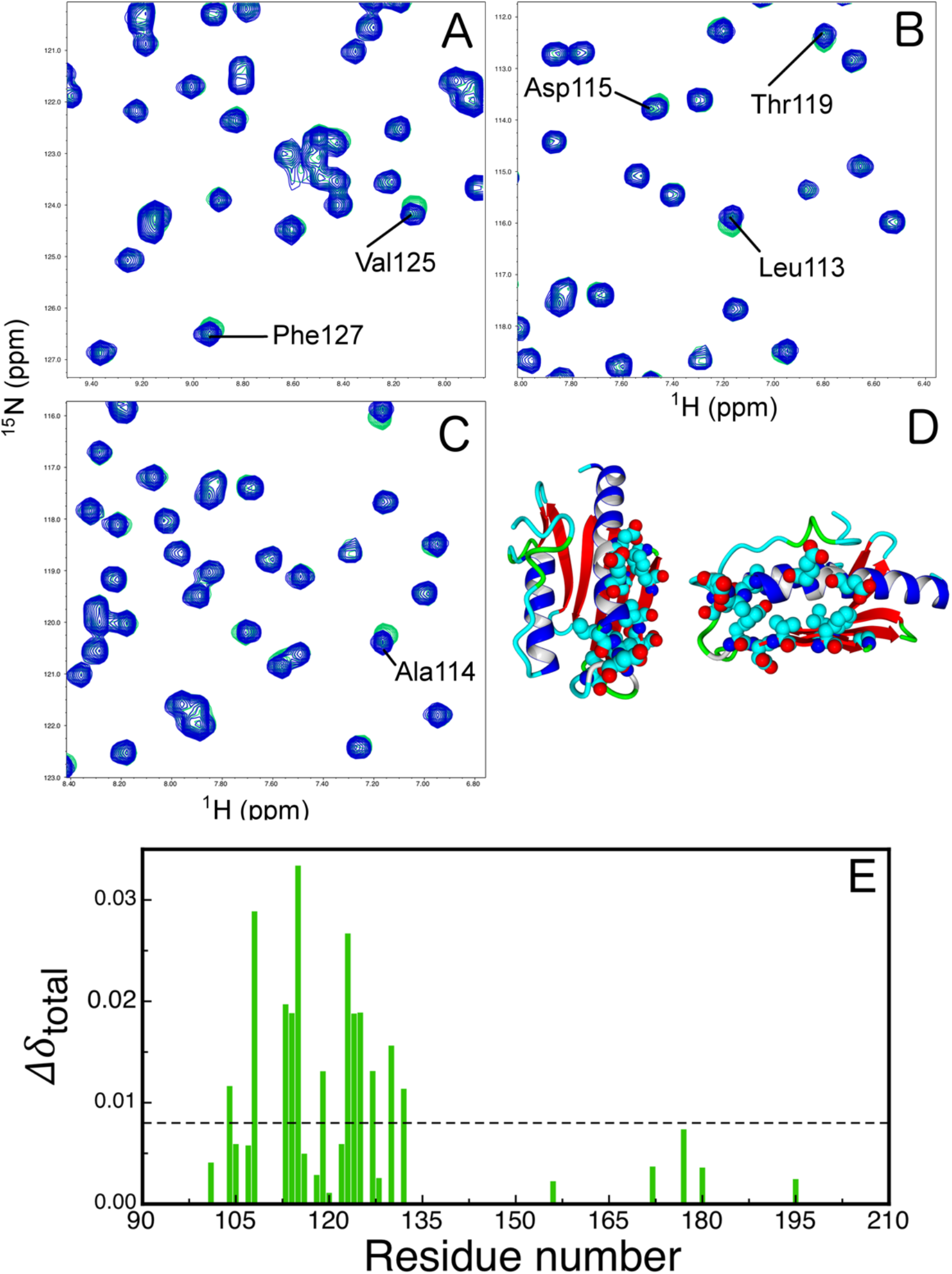
Characterization of Affi_224-frataxin Interaction by NMR. Selected regions of the ^1^N-^15^N HSQC spectra corresponding to ^15^N-labeled human frataxin (100 *μ*M) in the absence (blue) or in the presence (green) of Affi_224 (A, B, C). In (D), residues exhibiting high chemical shift perturbation are colored by element (two different views are shown). The CSP values calculated using Eqation 1 are plotted in (E). The dashed line indicates the cut off (two standard deviation over the average value) calculated as in reference [41]. Buffer was 20 mM Tris-HCl, 150mM NaCl, pH 7.0. Affi_224 was previously reduced by incubation with 1 mM TCEP. Affi_224 concentration was 200 *μ*M.

**Figure 7.**
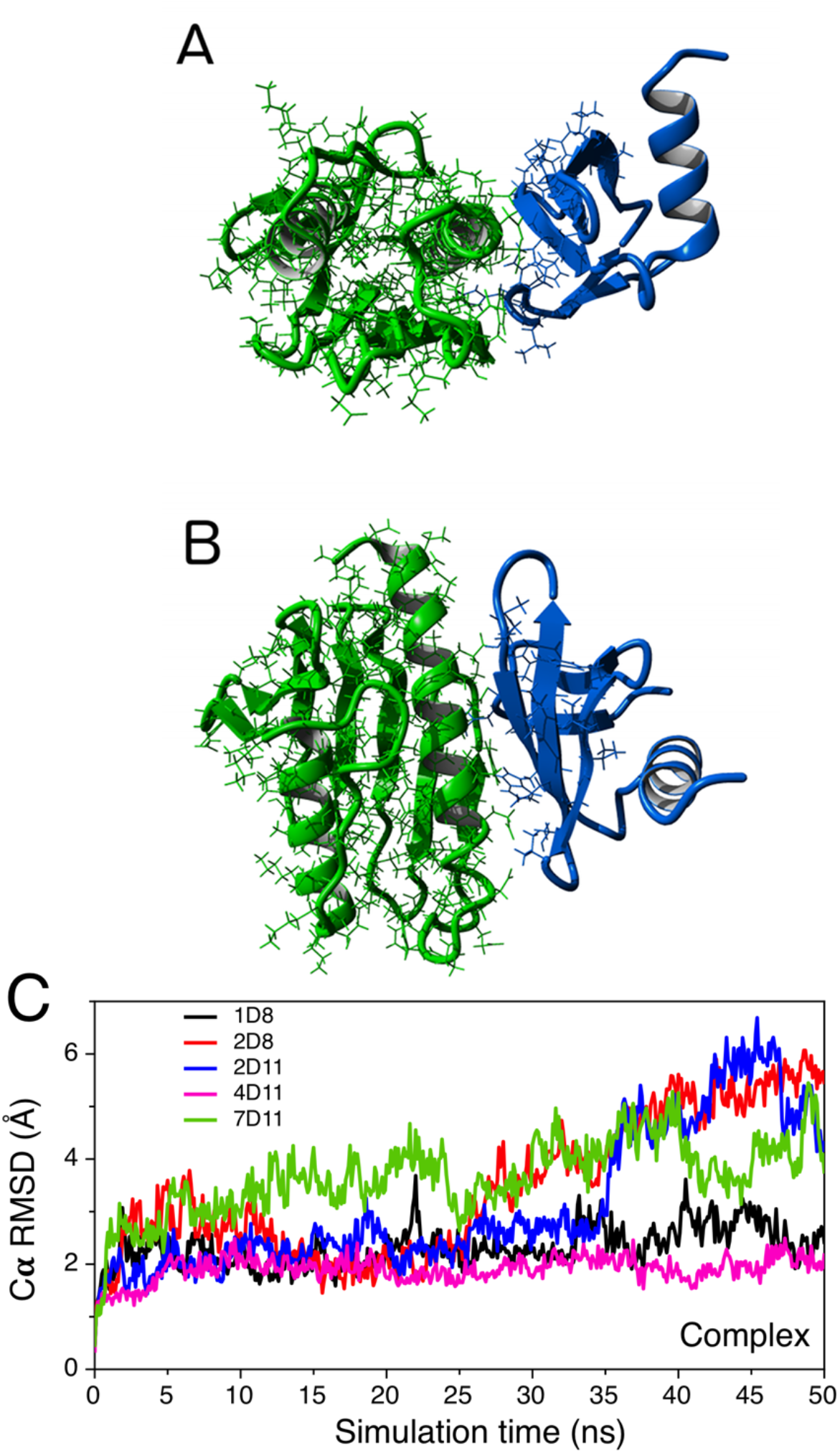
The Calculated Affi_224-Frataxin Complex. The molecular model corresponding to the hypothetical structure of the 1d8 Affi_224-frataxin complex is plotted; in (A) and (B) there are two different views of the complex. Side chains of affitin that were flexible during calculations are in blue sticks. Affi_224 and frataxin are in blue and green, respectively. (C) Molecular dynamics simulations of the hypothetical complexes. The alpha carbon RMSD values corresponding to the complexes calculated along the simulations of complexes 1D8, 2D8, 2D11, 4D11 and 7D11 (black, red and blue, magenta and green, respectively).

### Computational Characterization of Affi_224 / Frataxin Interaction

To explore the interaction mode between Affi_224 and frataxin at the atomic level, we performed global docking calculations using VINA [44] and the visual interface implemented in a Yasara Structure [45, 46]. For these calculations, a molecular model of Affi_224 was generated using the Swiss–model [47] and PDB ID, whereas 1SAP was the template (74.2% of identity). Next, all atom molecular dynamics simulation was carried out for affitin and frataxin to globally relax the conformation of the proteins (separated proteins, 50 ns independent runs), prior to docking experiments. The last snapshots were used for docking simulations. The residues that were the randomized positions during affitin selection (**Table S1**) were treated as flexible. On the other hand, the backbone and the rest of the side chains of both Affi_224 and frataxin were fixed during the simulations.

The analysis of the docking energies suggested that Affi_224 would interact with the frataxin acidic ridge (hypothetical model named 1d8, **Figs. 7 and S6A**). Even though this result is in complete agreement with the evidence provided by NMR experiments, we cannot exclude other possibilities because the difference in energy with other docking models is little. In **Table S1**, data concerning to the ten best models are shown. As expected, the interaction surface of the best of the complexes includes the majority of the residues that were considered flexible during the process. Frataxin–affitin contacting surfaces in those complexes that are similar in energy also involve several residues treated as flexible. However, in the 1d11 and 5d11 complexes, Affi_224 interacts by means of a completely different surface (**Figures S6 I and J** and **Table S1**).

To carry out a further computational evaluation of the Affi_224–frataxin interaction, we carried out short (50 ns) molecular dynamics simulations of the complexes using a Yasara Structure (**Figs. 7C** and **S7**). The instability of some of the Affi_224– frataxin complexes were confirmed by the increase of the RMSD values along the simulations (**Fig. 7C**), and the anomalous increase of the RMSF values (**Fig. S7A** and **B**) compared to the RMSFs measured for the molecular dynamics of more stable complexes or independent subunits. This analysis showed that the hypothetical complex 1d8 might be one of the more stable; in addition, the third complex 4d11, as judged by binding energy, might also be significantly stable.

### Frataxin-Affi_224 Interaction in the Cell Context

Whether or not this interaction occurs in complex environments is a key question. To investigate this issue, we expressed Affi_224 in a mammalian HEK-293T cell line. Cells were transfected with a pCDNA3.1_MTS-Affi_224 vector that codified Affi_224 preceded by the mitochondrial transit signal (MTS) from the citrate synthase enzyme for the mitochondrial matrix localization. Cell viability was evaluated after 48 h of transfection and there were no significant differences between transfection control (pCDNA 3.1 empty vector) and affitin transfected cells (not shown). Protein expression was evaluated by Western blotting using an anti RGSHis6 monoclonal antibody showing that Affi_224 was produced in significant fashion in this cell system (**Fig. 8A**).

**Figure 8.**
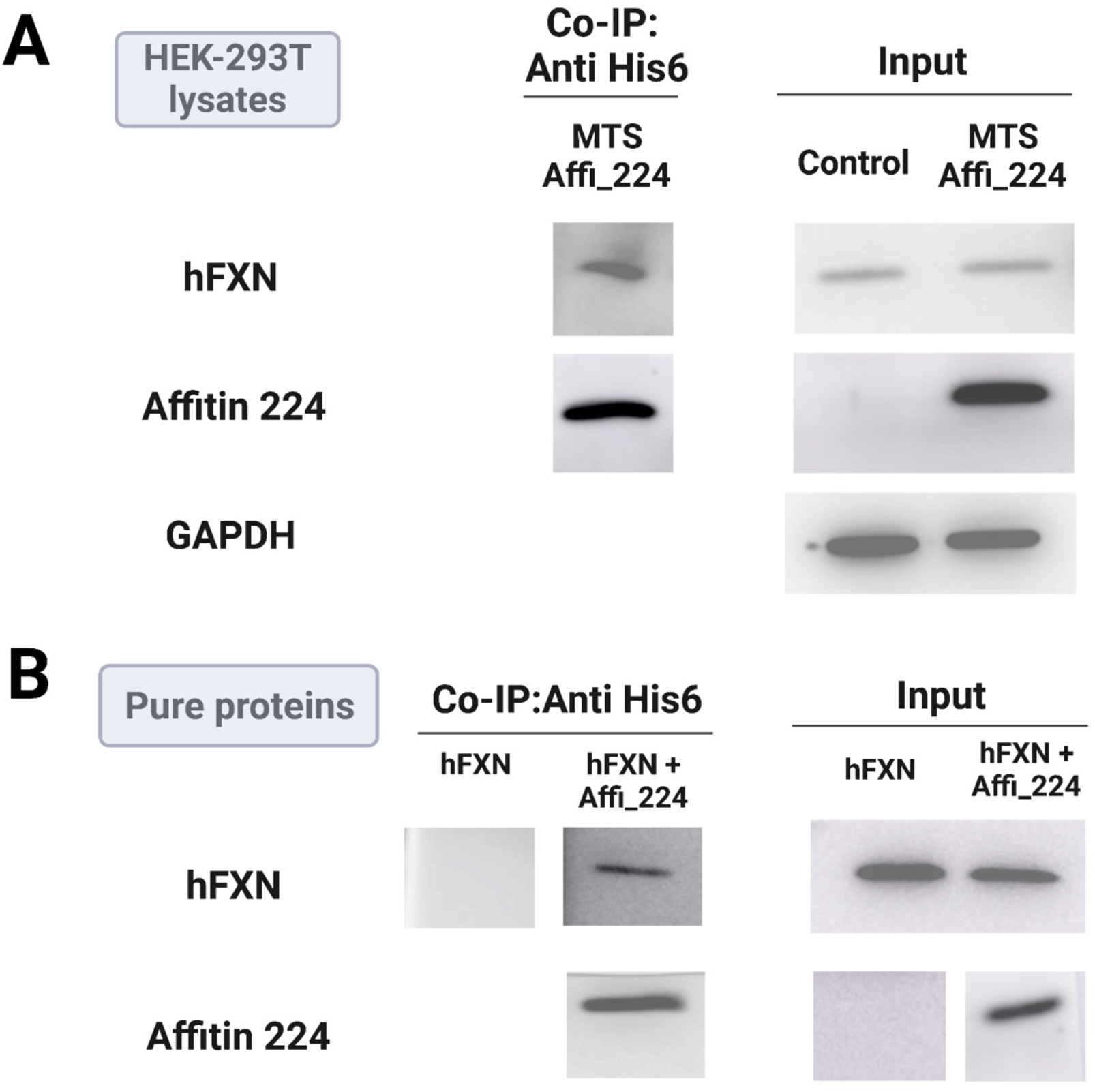
Co-immunoprecipitation of Frataxin and Affi_224. (A) Endogenous human frataxin was co-immunoprecipitated with an anti 6xHis mAB (Thermo Fisher). 250 μg of lysates from HEK-293T cells transfected with pcDNA3.1_MTS_Affi_224 or empty pcDNA3.1 were incubated with 2 μg of 6xHis mAB and the subjected to immunoprecipitation with G protein bound to magnetic beads (BioRad). (B) A similar experiment was also performed, albeit carried out with purified proteins using recombinant human frataxin 80-210 and Affi_224.

To set up the co-immunoprecipitation experiments, we first investigated the interaction using purified proteins (**Fig. 8B**). Incubation between frataxin and Affi_224 was carried out under standard conditions (1h at 25 °C in bufferTBS-0.05% Tween-20). After that, the anti His-tag monoclonal antibody was added (this antibody recognizes the His-tag in Affi_224) and immunoprecipitation was performed by means of protein G (using magnetic beads).

Western blotting was revealed using an anti-frataxin monoclonal antibody. Remarkably, the mAB anti frataxin recognized the FXN81-201 variant much better than the FXN90-210 variant (lane 3 vs 4). The same protocol was carried out to study the interaction in the cell environments by immunoprecipitation after cell disruption. Affi_224 expressed in HEK-293T cells with the mitochondrial targeting sequence of citrate synthase was co-immunoprecipitated with the endogenous frataxin.

### Affi_224 Interaction Modulates of Cys Desulfurase Activation

Affi_224 was able to bind to frataxin in a clear-cut manner. However, affitin interaction through the frataxin acidic ridge may inhibit frataxin binding to the supercomplex for Fe-S cluster assembly. In particular, frataxin interacts via these acidic residues with a positively charged stretch of one of the subunits of the desulfurase enzyme dimer (PDB ID: 6NZU) [22].

To evaluate this possibility, L-Cys desulfurase activity of NFS1 was tested in the absence or in the presence of Affi_224 (**Fig. 9**). The addition of Affi_224 to the reaction did not produce significant inhibition, suggesting that the interaction does not inhibit NFS1-frataxin binding.

**Figure 9.**
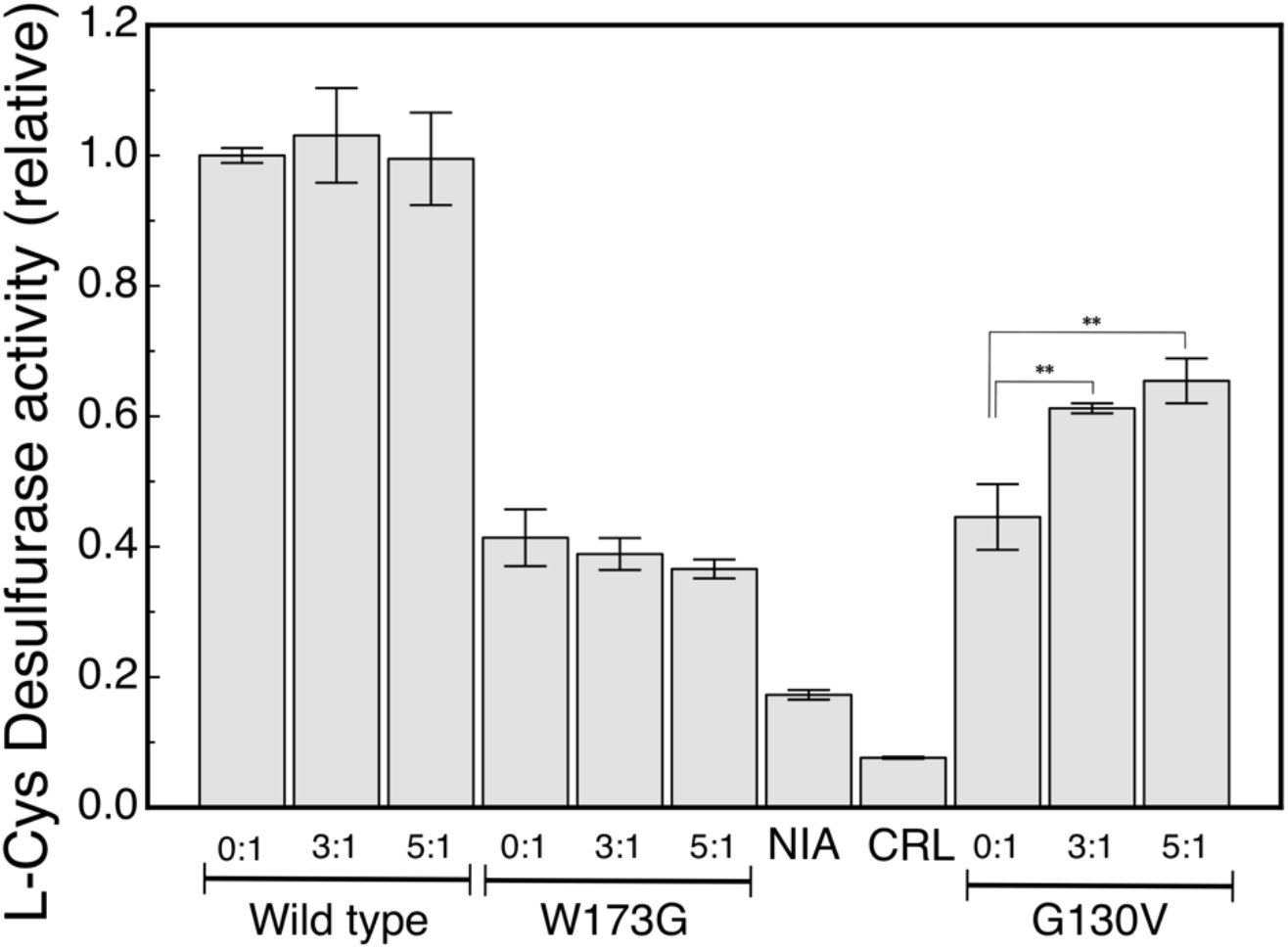
Modulation of Cys Desulfurase Activity by Affi_224. Activation is exerted by a wild–type frataxin, the W173G or G130V variant, in the presence or absence of Affi_224. Affitin was in two different concentrations, 9.0 and 15 μM (3 and 5 times the concentration of frataxin in molar ratio: 3:1 and 3:5, respectively). Reactions contained 1.0 μM NFS1, 1.0 μM ACP-ISD11, 3.0 μM ISCU and 3.0 μM frataxin. Samples were supplemented with 10 μM PLP, 2 mM DTT, and 1.0 μM FeSO4 (final concentrations). The reaction buffer was 50 mM Tris-HCl and 200 mM NaCl, pH 8.0, and reactions were started by the addition of 1 mM cysteine. Samples were incubated at room temperature for 30 min. H_2_S production was stopped by the addition of 50 μL of 20 mM N,N-dimethyl p-phenylenediamine in 7.2 M HCl and 50 μL of 30 mM FeCl_3_ (prepared in 1.2 M HCl). Under these conditions, the production of methylene blue took 20 min. After that, samples were centrifuged for 5 min at 12000g, and the supernatant was separated. The absorbance at 670 nm was measured. Control reactions without frataxin and ISCU (NIA), or without proteins (CRL), were performed. The difference was considered statistically significant when p<0.003 (**, Tukey’s multiple comparison test).

On the other hand, we studied the effect of the addition of Affi_224 to the G130V and W173G frataxin variants. In the case of the latter, we did not observe an improvement of the activation. The activity of the W173G variant was very low both in the absence and in the presence of Affi_224. However, the G130V variant exhibited a small but reproducible and significant increase of L-Cys-desulfurase activation, suggesting that not only Affi_224 would not be an inhibitor of wild–type frataxin but it may also positively modulate the activity of specific FRDA frataxin variants. More experiments should be performed to study the molecular basis of this activation and Affi_224 effects on physiological Fe-S cluster assembly reaction.

As interaction in the hypothetical complex 1d8 (and detected by NMR) may be incompatible with frataxin binding to NFS1, an alternative hypothesis is that Affi_224 interaction is not preserved when frataxin binds to the supercomplex. This fact might be compatible with a rapid equilibrium of binding for Affi_224–frataxin interaction and a slow dissociation of frataxin from the supercomplex. An interesting issue that has yet to be studied is whether Affi_224 is able to stabilize the active form of other unstable frataxin variants.

## Conclusions

We were able to obtain anti frataxin affitins by applying the ribosome display selection strategy. The selected affitins showed a significant propensity to form dimers and some of them contained Cys residues that can form disulfide bonds. They interacted with chromatographic media in SEC-HPLC experiments. In particular, Affi_224 was only marginally stable. These issues might make the use of Affi_224 highly challenging. However, Affi_224 showed a *K*_D_ value in the nanomolar range and, most likely, it binds to frataxin through interaction involving the frataxin acidic ridge, as judged by the analysis of CSP and computational simulations.

Furthermore, it is worthy of note that Affi_224 was able to increase Cys NFS1 desulfurase activation exerted by the FRDA frataxin variant G130V. Affi_224 did not alter the activation exerted by the wild–type variant of frataxin, suggesting that it does not inhibit frataxin binding to the supercomplex. On the other hand, Affi_224 was not able to increase the activity of the highly unstable FRDA frataxin variant W173G.

Our results suggest quaternary addition may be a new tool to modulate frataxin function *in vivo*. Nevertheless, more functional experiments under physiological conditions should be carried out to evaluate Affi_224 effectiveness in FRDA cell models.

## Acknowledgments

We especially thank FARA, CONICET and Universidad de Buenos Aires for financial support. We thank F.J. Pérez De Berti from Centro de Investigación en Hidratos de Carbono (CONICET) for his help with CD measurements. We would like to especially thank Dr. Agustin Correa from Instituto Pasteur Montevideo, Uruguay, for his support during the selection of affitins.

## Author Contributions

MFP, GH and JS planned the study, performed the experiments and wrote the paper.

## UniProt Accession IDs

Q9HD34: human LYR motif-containing protein 4 (ISD11); O14561: human mitochondrial acyl carrier protein (ACP); Q9Y697: human cysteine desulfurase (NFS1); Q9H1K1: human iron–sulfur cluster assembly enzyme (ISCU); Q16595: human frataxin (FXN).

## Abbreviations

ACP: acyl carrier protein
CD: circular dichroism
CTR: C-terminal region
DLS: dynamic light scattering
ELISA: enzyme-linked immunosorbent assay
Fe-S: iron–sulfur
FRDA: Friedreich’s Ataxia
FXN: frataxin
HPLC: high-performance liquid chromatography
ISCU: iron-sulfur cluster assembly enzyme
ISD11: NFS1 interacting protein
NFS1: mitochondrial cysteine desulfurase enzyme
NMR: nuclear magnetic resonance
PAGE: polyacrylamide gel electrophoresis
PDB: Protein Data Bank
Sac7D: a small DNA binding protein from the hyperthermophilic archaeon *Sulfolobus acidocaldarius*
SDS: sodium dodecyl sulfate
SEC: size exclusion chromatography
SPR: surface plasmon resonance

