## Supplementary Information for "Selection of Synthetic Proteins to Modulate the Human Frataxin Function"

[1] Instituto de Biociencias, Biotecnología y Biología Traslacional (iB3). Departamento de Fisiología y Biología Molecular y Celular, Facultad de Ciencias Exactas y Naturales, Universidad de Buenos Aires, Intendente Güiraldes 2160, Ciudad Autónoma de Buenos Aires, Argentina.
[2] Consejo Nacional de Investigaciones Científicas y Técnicas, Av. Rivadavia 1917 (C1033AAJ), Ciudad Autónoma de Buenos Aires, Argentina.
[3] Fundación Instituto Leloir, IIBBA-CONICET, Av. Patricias Argentinas 435, C1405BWE, Buenos Aires, Argentina.
[4] Departamento de Química Inorgánica, Analítica y Química Física, Facultad de Ciencias Exactas y Naturales, Universidad de Buenos Aires. Instituto de Química Física de los Materiales, Medio Ambiente y Energía (INQUIMAE CONICET), C1428EGA, Buenos Aires, Argentina

**Running Title:** Frataxin-Affitin Interaction

**Keywords:** conformational stability, protein–protein interaction, iron–sulfur cluster assembly

**Abbreviations:** ACP, acyl carrier protein; CD, circular dichroism; CTR, C-terminal region; DLS, dynamic light scattering; ELISA, enzyme-linked immunosorbent assay; Fe-S, iron–sulfur; FRDA, Friedreich's Ataxia; FXN, frataxin; HPLC, high-performance liquid chromatography; ISCU, iron-sulfur cluster assembly enzyme; ISD11, NFS1 interacting protein; NFS1, mitochondrial cysteine desulfurase enzyme; NMR, nuclear magnetic resonance; PAGE, polyacrylamide gel electrophoresis; PDB, Protein Data Bank; Sac7D, a small DNA binding protein from the hyperthermophilic archaeon *Sulfolobus acidocaldarius*; SDS, sodium dodecyl sulfate; SEC, size exclusion chromatography; SPR, surface plasmon resonance.

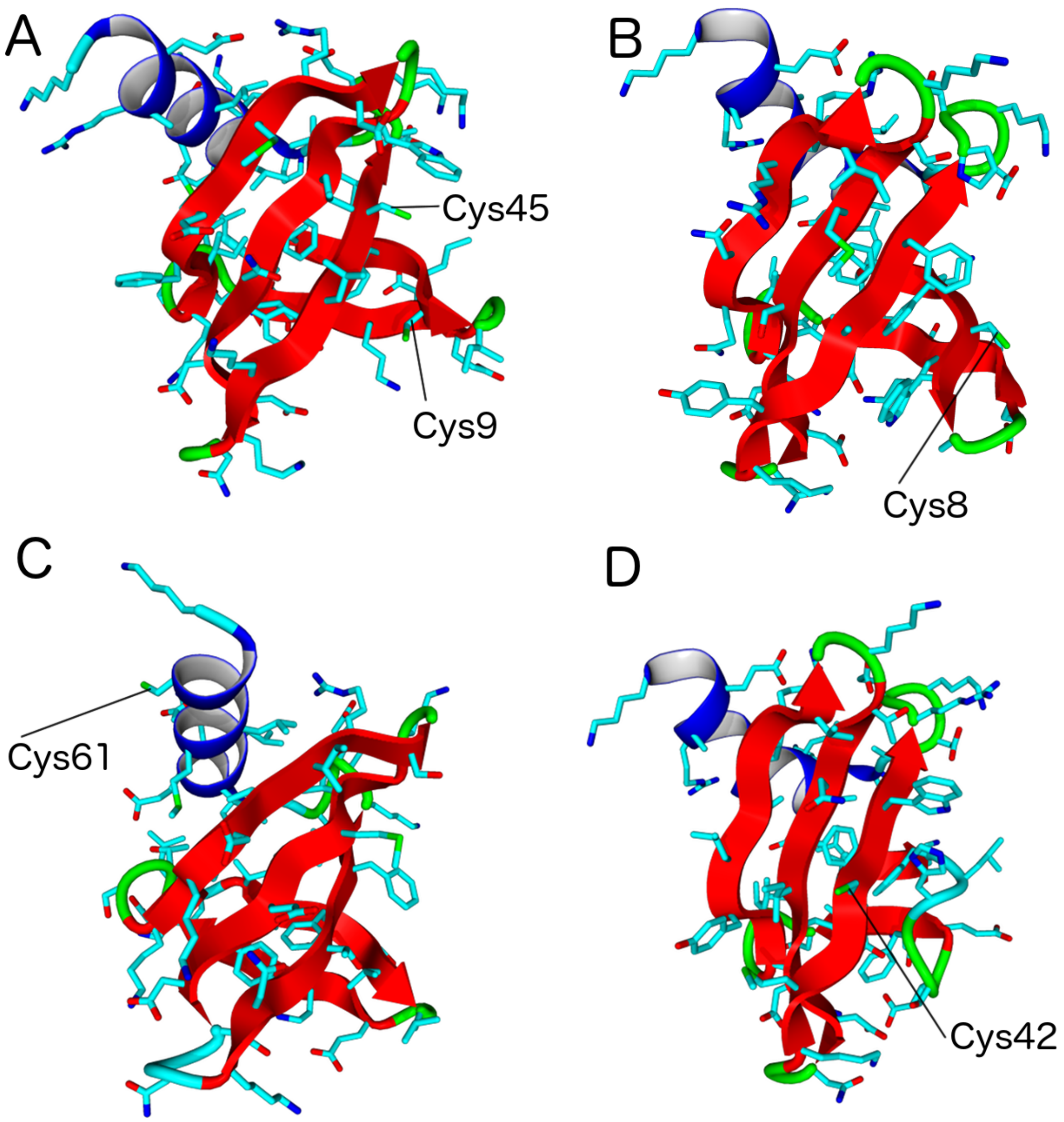


**Figure S1. Affitin Structure Models.** (A) Affi_37. (B) Affi_60. (C) Affi_186. (D) Affi_224. Cys residues are indicated and models were prepared with the Swiss Model. Solvent–accessible surface areas (SASA): Cys9 and Cys45: 34 and 28%, respectively; Cys8: 71.8%; Cys61: 81.9%; Cys42: 66.9%. SASA was calculated using GETAREA http://curie.utmb.edu/getarea.html. Fraczkiewicz, R. and Braun, W. (1998) "Exact and Efficient Analytical Calculation of the Accessible Surface Areas and Their Gradients for Macromolecules", J. Comp. Chem., 19, 319-333.

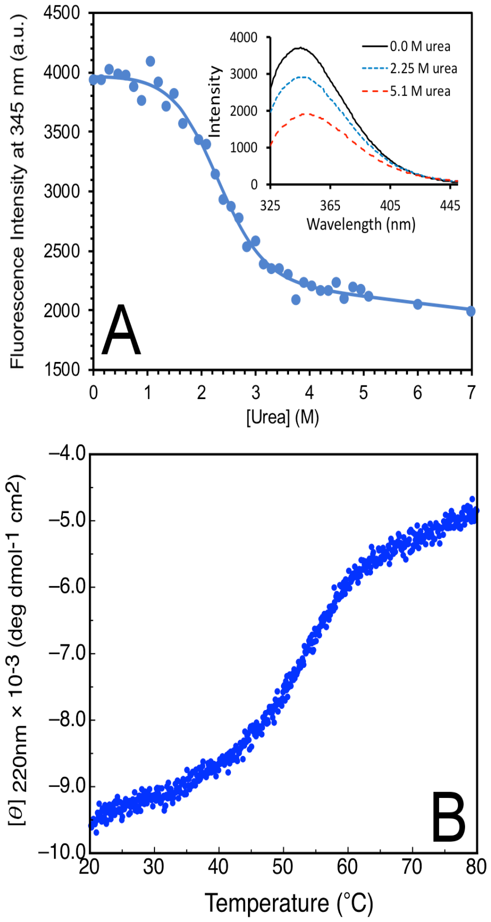


**Figure S2. Conformational Stability of Affi_224.** (A) Urea–induced unfolding of Affi_224. Equilibrium unfolding was monitored by tryptophan fluorescence. Excitation was carried out at 295 nm and a bandwidth of 5 nm was used for excitation and emission. For each urea concentration, two spectra were recorded and averaged. Dependence of the intensity at 345 nm with urea concentration was plotted. The inset shows fluorescence spectra at three different urea concentrations. Buffer was 20 mM Tris-HCl, 150mM NaCl and pH 7.0. (B) Temperature–induced unfolding of Affi_224 in 25 mM sodium phosphate, 100mM NaCl, pH 7.0.

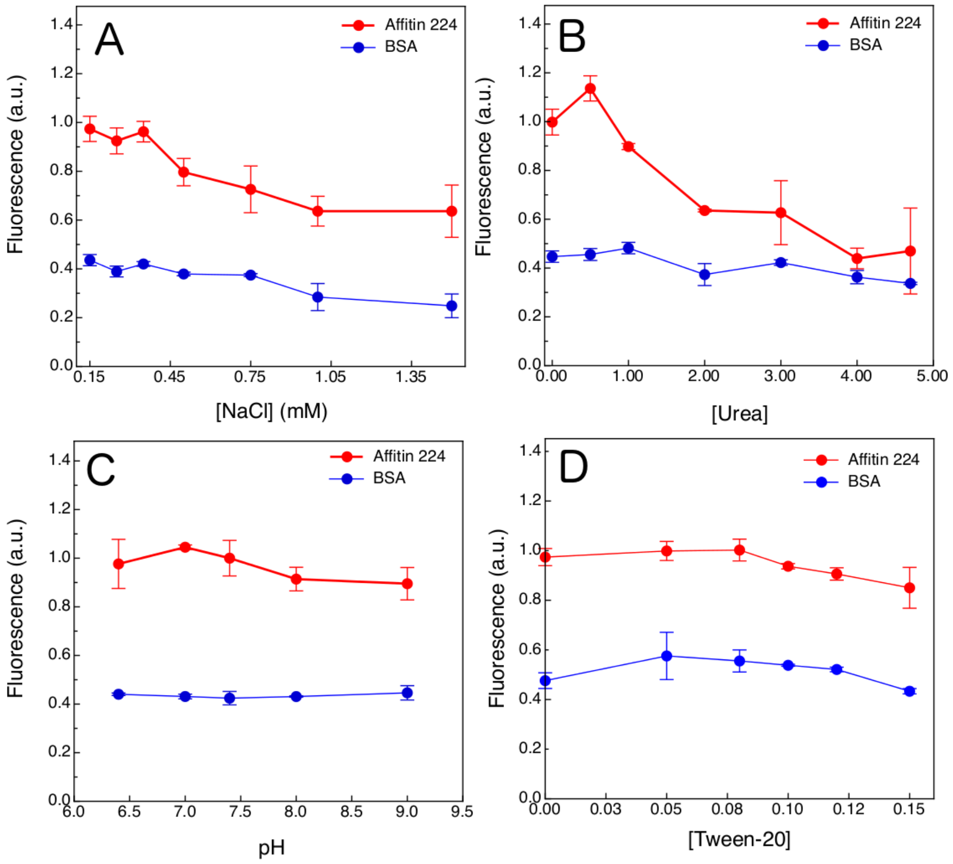


**Figure S3. Perturbation of Affi_224-frataxin Interaction Monitored by Fluorescence.** Affi_224 was bound to the multi–well plates blocked by the addition of BSA. Frataxin labeled with Texas Red was added in the presence or absence of NaCl (A) or urea (B). Also, the proteins were incubated at different pH values (C). Different concentrations of Tween-20 were also tested (D). Control wells containing only BSA were assayed for detection of unspecific interactions of labeled frataxin (blue). All measurements were carried out under the same experimental conditions: 20 mM Tris-HCl, 150mM NaCl and pH 7.0.

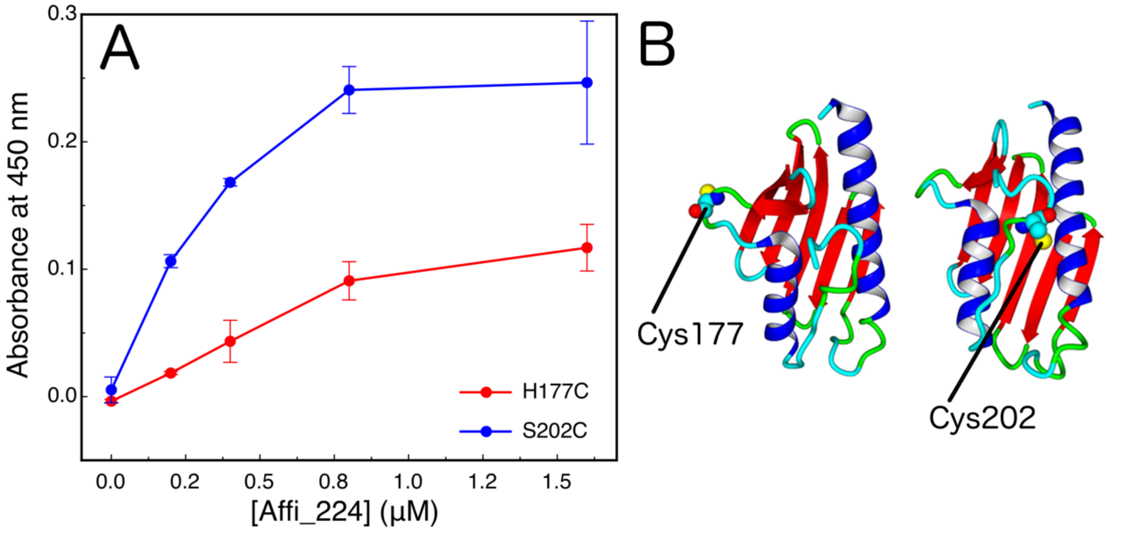


**Figure S4. Interaction between Frataxin and Affi_224 Monitored by ELISA.** (A) Biotinylated frataxin variants H177C and S202C were bound to the multi–well neutravidin–coated plates; after that, the surfaces were blocked by the addition of BSA. Increasing concentrations of Affi_224 were added to different wells. The detection of the interaction was performed by an RGS His6 HRP conjugate that detects the RGS His6 motif present in the affitin N-terminal. A dilution of the antibody at 1:4000 in TBS-Tween 0,1% was used for ELISA detection. The horseradish–peroxidase substrate was TMB. Absorbance at 450 nm was read after the addition of 4N sulfuric acid. The location of the Cys177 and Cys 202 residues are shown in (B).

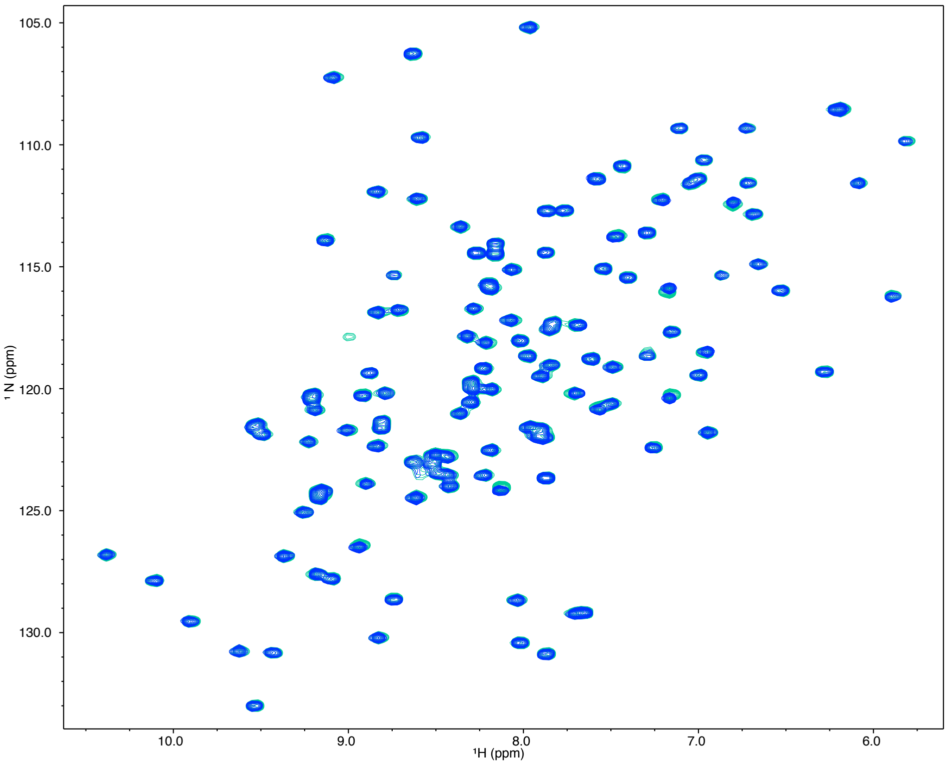


**Figure S5. $^{1}$N-$^{15}$N HSQC Spectra Corresponding to 15N-labeled Frataxin (100 $\mu$M) in the Absence (blue) or in the Presence (green) of Affi_224.** Buffer was 20 mM Tris-HCl, 150mM NaCl, pH 7.0. Affi_224 was previously reduced by incubation with 1 mM TCEP. Affi_224 concentration was 200 $\mu$M.

**Table S1. Global Docking Calculations[2] and the Analysis between Affi_224 and Frataxin Interaction.**

| Docking | Predicted Binding Energy (kcal/mol) | Predicted Dissociation Constant (pM) | Contacting Residues[1] |
|---|---|---|---|
| **1d8** | 14.9 | 11.8 | Phe7, **Leu8**, **His 9**, **Phe10**, **Val26**, Lys28, **Thr29**, **Asn31**, Tyr34, Lys39, **Met40**, Gly41, **Cys42**, **Trp44** |
| **2d11** | 14.5 | 23.2 | Phe7, **His9**, **Tyr21**, **Val23**, **Gly24**, Arg25, **Val26**, Lys28, **Thr29**, **Asn31**, **Ile33**, Tyr34, **Met40**, Gly41, **Cys42**, Gly43, **Trp44** |
| **4d11** | 14.0 | 53.9 | **Tyr21**, **Leu22**, **Val23**, **Gly24**, Arg25, **Val26**, **Asn31**, **Ile33**, Tyr34, Asp35, **Met40**, **Cys42**, Gly43, Leu58, Arg60, Ala61, GLu62, Gly63, Lys64 |
| **7d11** | 14.1 | 50.1 | **Tyr21**, **Leu22**, **Val23**, **Gly24**, Arg25, **Val26**, **Asn31**, Phe32, **Ile33**, **Met40**, **Cys42**, Gly43, **Trp44**, Leu58, Ala61, GLu62, |
| **2d8** | 13.9 | 68.1 | **Tyr21**, **Leu22**, **Val23**, **Gly24**, Arg25, **Val26**, **Asn31**, **Ile33**, Lys39, **Met40**, Gly41, **Cys42**, **Trp44**, Leu58, Ala61, Glu62 |
| **3d11** | 13.8 | 82.4 | Phe7, **His9**, **Tyr21**, **Val26**, Lys28, **Thr29**, **Asn31**, **Ile33**, Tyr34, Lys39, **Met40**, Gly41, **Cys42**, Gly43, **Trp44** |
| **8d11** | 13.4 | 155.5 | Phe7, **Leu8**, **His9**, **Val26**, Gly27, Lys28, **Thr29**, **Trp44**, Arg46, Glu47, Lys48 |
| **9d11** | 13.0 | 303.8 | Phe7, **His9**, **Val26**, Gly27, Lys28, **Thr29**, **Asn31**, **Ile33**, Lys39, **Met40**, Gly41, **Cys42**, Gly43, **Trp44** |
| **1d11** | 12.8 | 402.1 | Arg25, Lys28, Glu47, Lys48, Lys52, Leu55, Asp56, Leu58, Ala59, Arg60, Glu62, Gly63, Lys64 |
| **5d11** | 12.1 | 1314.8 | Val3, Lys4, ASp16, Thr17, Ser18, Lys19, Ile20, **Tyr21**, Asp35, Asp36, Asn37, Glu53, Met57, Arg60, Gly63, Lys64, |

[1] Residues of Affi_224 that interact with frataxin in this complex.

[2] Residues that were flexible during the docking calculations: Leu8, His9, Phe10, Ser11, Tyr21, Leu22, Val23, Gly24, Val26, Thr29, Asn31, Ile33, Met40, Cys42 and Trp44.

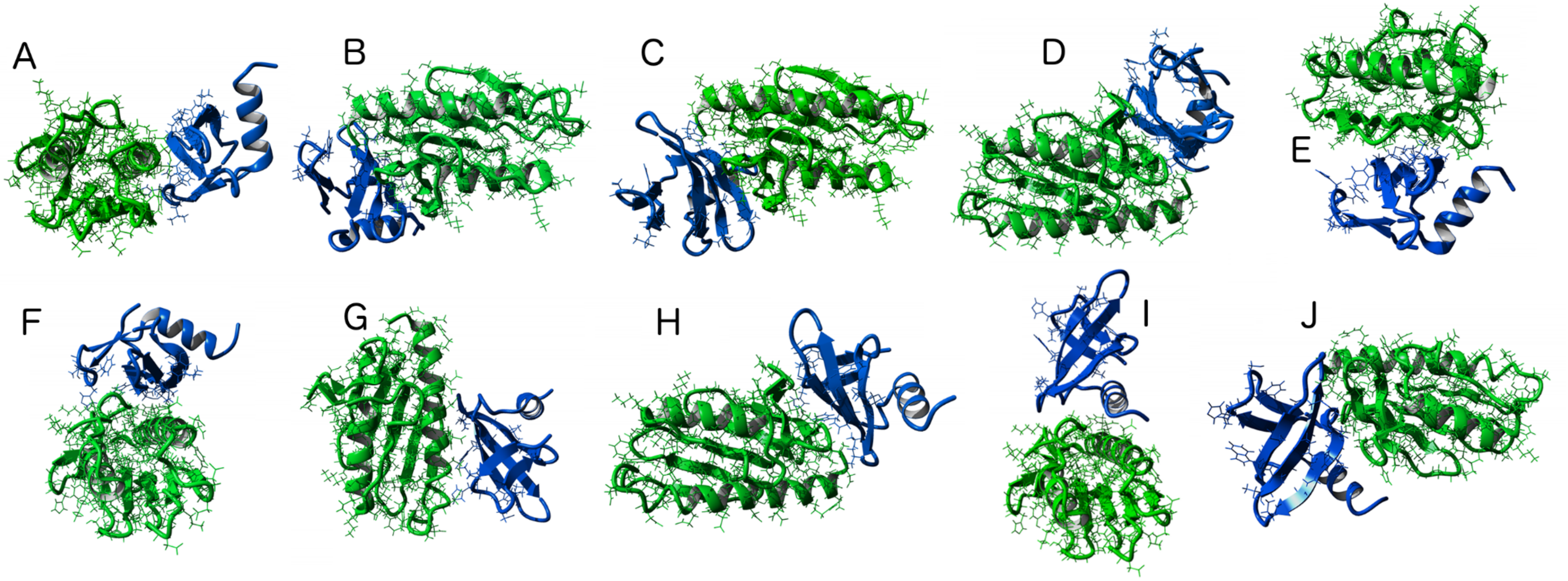


**Figure S6. Calculated Affi_224-frataxin Complexes.** Molecular models corresponding to the hypothetical structures of Affi_224-frataxin complex are plotted: 1d8 (A), 2d11 (B), 4d11 (C), 7d11 (D), 2d8 (E), 3d11 (F), 8d11 (G), 9d11 (H), 1d11 (I), 5d11 (J).

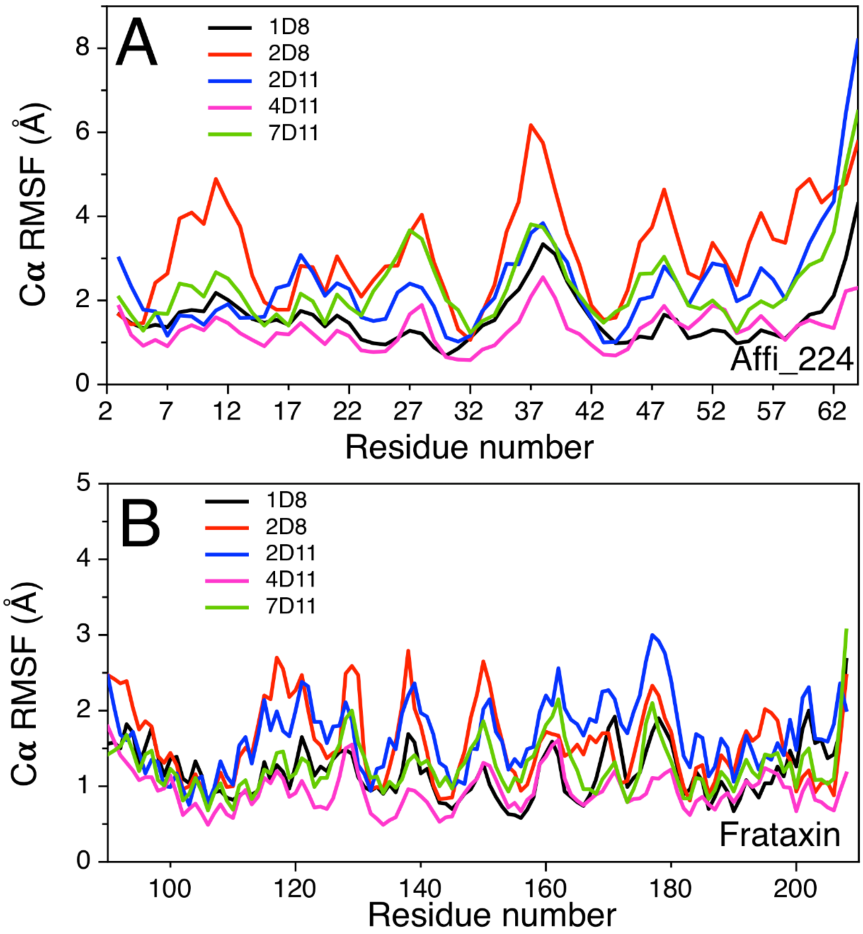


**Figure S7. Molecular Dynamics Simulations of the Hypothetical Complexes.** The alpha carbon RMSFs calculated using the trajectories of the respective complexes and the alpha carbon RMSD values corresponding to the complex are shown in (A) and (B), corresponding to Affi_224 and frataxin, respectively, 1D8, 2D8, 2D11, 4D11 and 7D11 are in black, red and blue, magenta and green, respectively. The RMSD along the simulations are shown in Figure 7C, main text.
